# Dynamic phytomeric growth contributes to local adaptation in barley

**DOI:** 10.1101/2023.06.02.543309

**Authors:** Yongyu Huang, Andreas Maurer, Ricardo F. H. Giehl, Shuangshuang Zhao, Guy Golan, Venkatasubbu Thirulogachandar, Guoliang Li, Yusheng Zhao, Corinna Trautewig, Axel Himmelbach, Andreas Börner, Murukarthick Jayakodi, Nils Stein, Martin Mascher, Klaus Pillen, Thorsten Schnurbusch

## Abstract

Vascular plants segment their body axis with iterative nodes of lateral branches and internodes. Appropriate node initiation and internode elongation are fundamental to plant fitness and crop yield formation; but how they are spatiotemporally coordinated remains elusive. We show that in barley (*Hordeum vulgare* L.), selections under domestication have extended the apical meristematic phase to promote node initiation, but constrained subsequent internode elongation. In both vegetative and reproductive axes, internode elongation displays a dynamic proximal – distal gradient, and among subpopulations of domesticated barleys at the global range, node initiation and proximal internode elongation are associated with latitudinal and longitudinal gradients, respectively. Genetic and functional analysis suggest that, in addition to their converging roles in node initiation, flowering time genes are repurposed to specify the dynamic internode elongation. Our study provides an integrated view of barley node initiation and internode elongation, and suggests that plant architecture has to be recognized as dynamic phytomeric units in the context of crop evolution.

## Introduction

Plant architecture is the outcome of several successive developmental processes that can be classified into two sequential and coordinated morphogenetic events: organogenesis and extension^1^. Organogenesis stems from the indeterminate, self-renewing meristems (stem cells) that give rise to different types of lateral organs (e.g., leaves and flowers) and axillary bud(s), plus the subtending internodes. These apex-derived organs form a functional unit, called phytomer^2^ that iterates and extends itself several rounds until the apex abscises/aborts (indeterminate growth) or terminates into a specialized structure (determinate growth). Consequently, a plant’s architecture is a mosaic arrangement of different phytomers. Despite an individual’s genetic uniformity, these mosaic phytomers can adopt various shapes in response to exogenous environmental constraints by, for example, altering the meristematic determinacy or internode elongation^3, 4^. Studying plant architecture is therefore of fundamental importance to understand plant developmental biology and environmental adaptation.

Barley (*Hordeum vulgare* L.) is a diploid inbreeding species and can be considered as a model of the *Triticeae* crops, including also wheat (*Triticum* sp.) and rye (*Secale cereale* L.). Plant architecture of barley consists of a two-ranked (distichous) arrangements of both leaves and floral organs (spikelets) on the alternate sides of vegetative culms and the reproductive axis (rachis), respectively. In this context, the barley’s main body axis represents a simple and continuous segmentation of phytomers wherein both vegetative and reproductive organs co-exist at the opposite ends (Fig. 1a). As an economically important crop that is usually grown at a high- planting density where canopy shade is prevalent (e.g., ∼300 plants/m^2^ in Europe), genetic modification aiming at improving barley’s plant architecture may need to be implemented in the direction of high community uniformity with stable grain yield formation^5^. Yet, individual yield maximization may sometimes conflict with community performance^6, 7^, in particular, for less lodging-tolerant crops like barley^8^. Therefore, a more fundamental understanding of how barley plant architecture is controlled is of high agricultural relevance.

**Fig. 1:**
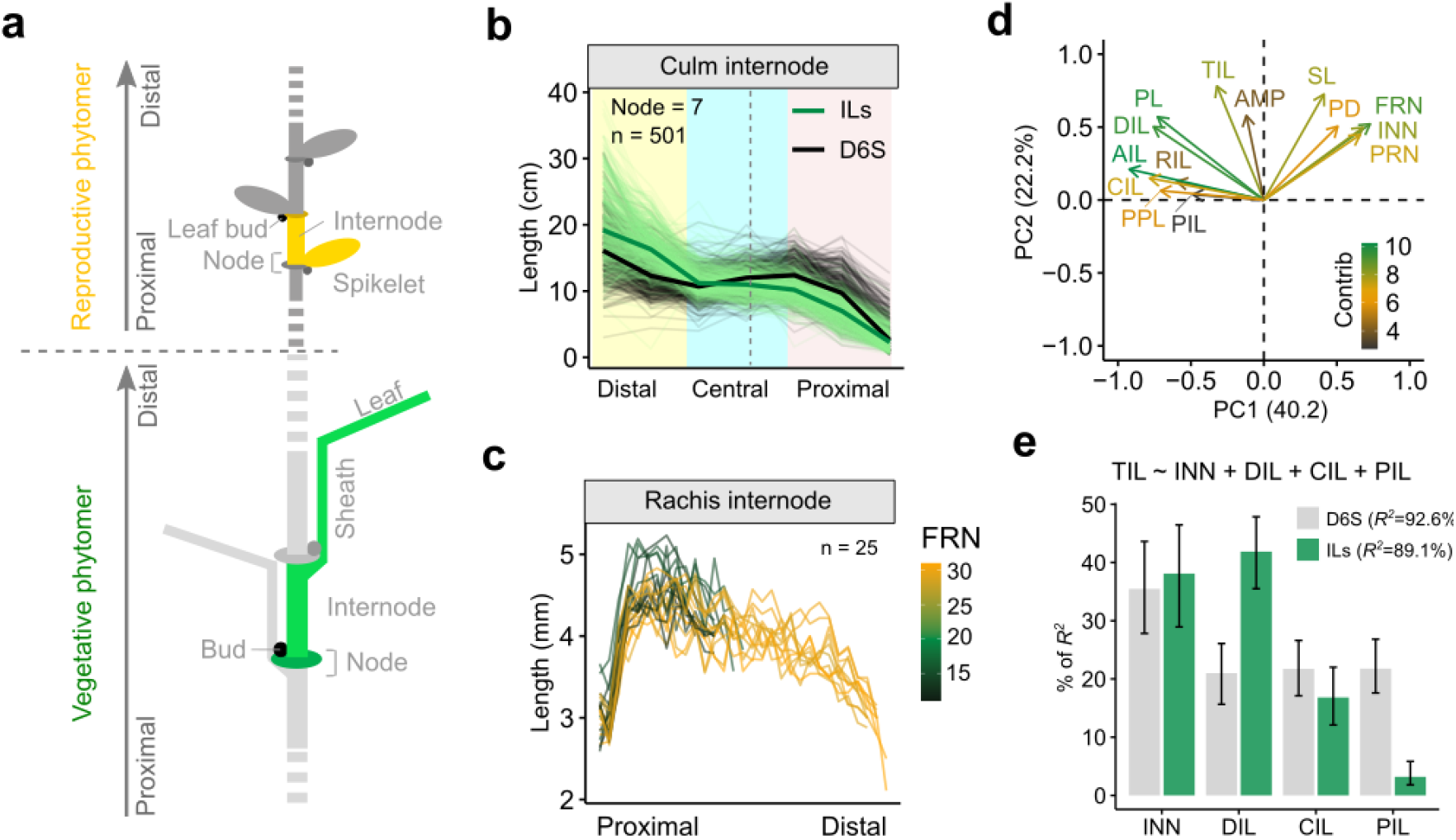
An oscillatory pattern of internode elongation and the node initiation – internode elongation relationship. **a.** Schematic representation of phytomeric structure from a barley vegetative culm and reproductive spike based on^2^. A vegetative culm phytomer and a reproductive spikelet phytomer is highlighted with green and orange colors, respectively. Drawings are not to scale. **b, c.** Pattern of internode elongation from culms (**b**) and rachises (**c**). Plants with a node number of 7 are shown in (**b**). **d.** Loading plot of PC1 and PC2 from the 14 phenotypes representing node initiation and internode elongation. Note that initiation and elongation related traits are largely separated based on the first two PCs. DIL, distal internode length; CIL, central internode length; PIL, proximal internode length; AIL, average internode length; PL, peduncle length; PPL, percentage of peduncle length; TIL, total culm internode length; AMP, culm length oscillatory amplitude; SL, spike length; PD, peduncle diameter; FRN, final rachis node number; INN, culm node number; PRN, potential rachis node number. **e**. Relative importance INN, DIL, CIL and PIL to total culm length (TIL). A multiple linear regression analysis is formulated on the top of the bar plot. Upper and lower bound of the error bars are 95% confidence intervals based on 1000 bootstrap replicates.

Signals orchestrating plant architecture appear to act on several phytohormones and their cross-talks, among which gibberellin (GA) has a fundamental and essential role for cell elongation^3, 9^. In particular, mutant alleles in the GA biosynthesis or signaling pathways have been demonstrated to improve grain yield potential during the ‘Green Revolution’ of wheat and rice (*Oryza sativa* L.) in the 1960s^10^. Other phytohormones such as brassinosteroids (BRs), jasmonate (JA) and ethylene were also found to modulate internode elongation via the GA pathway^11–15^. However, GA by itself is not essential for organogenesis^16^; instead, the florigen/antiflorigen, encoded by the *FLOWERING LOCUS T* (*FT*)/*TERMINAL FLOWER 1* (*TFL1*) family genes, are a key determinant of organ number by modifying the phase durations and expression of organ identity genes^17^. In barley, the activation of, e.g., *HvFT1* is modulated by several endogenous and exogenous cues and their cross-talks, such as circadian clock, photoperiod and temperature (i.e., vernalization)^18–22^. Mutations in any of these flowering time genes frequently result in changes of both vegetative and reproductive phytomeric iterations^23^, presumably via modulating meristematic determinacy^24^. In fact, floral induction is often coupled with an increase of GA concentration in the vegetative apex; consequently, many flowering plants elongate their internodes only after floral induction^25^, suggesting a cooperative action of the two hormonal systems [GA and (anti-) florigen] during morphogenesis. Still, how phytomer initiation and elongation are coordinated during morphogenesis remains less understood.

Here, we systemically investigated phytomer initiation and elongation by focusing on node number and internode length from the vegetative culms and reproductive spikes of barley (Supplementary Fig. S1). We used a series of wild barley (*Hordeum vulgare* subsp. *spontaneum*) introgression lines (ILs) in the background of cv. Barke (an elite two-rowed cultivar)^26^, as well as a diversity panel of cultivated barleys (spring-type, six-rowed, hereafter D6S) representing the three major global-wide subpopulations relative to the center of barley origin (i.e., Eastern, Western and Ethiopian clades)^27^ (Supplementary Table 1). The underlying assumption is that over thousands of years of barley evolution, the spread and fixation of beneficial alleles may have enabled a fine-tuning of barley plant architecture to better adapt to the local environments or high-density agricultural practices. Our results suggest that the barley plant (entity) is compartmentized into dynamic phytomeric units, with distinct flowering time genes functionally repurposed to determine their initiation and/or elongation. We provide evidence that proximal internode elongation, a previously underexplored functional trait, is associated with both plant adaptation and reproductive efficiency (spikelet survival).

## Results

### A more compact inflorescence architecture under domestication

We primarily focused on reproductive traits in an initial survey for phenotypic diversity in the 25 wild barley founder parents of the HEB-25 population^26^ (Supplementary Fig. 2a,b). We found that all inflorescence meristems from wild barley spikes were developmentally arrested and degenerated much earlier compared to cv. Barke; consequently, wild barleys had both lowered potential rachis node number (PRN) at the maximum yield potential stage^28^ and lowered final rachis node number (FRN) at the anthesis stage. Notably, spike length (SL) was largely unchanged, consequently, wild barleys had longer rachis internode length (RIL) than cv. Barke. We conclude that evolution under domestication may have overall extended the apical meristematic phase to promote node initiation, but compromised the subsequent internode elongation, resulting in a more compact inflorescence architecture. To further uncover the genetic underpinnings of these events, we selected four sub- families comprising 247 introgression lines (ILs) based on both the phenotypic and genotypic diversity of the wild barley gene pool (Supplementary Fig. 2c).

**Fig. 2:**
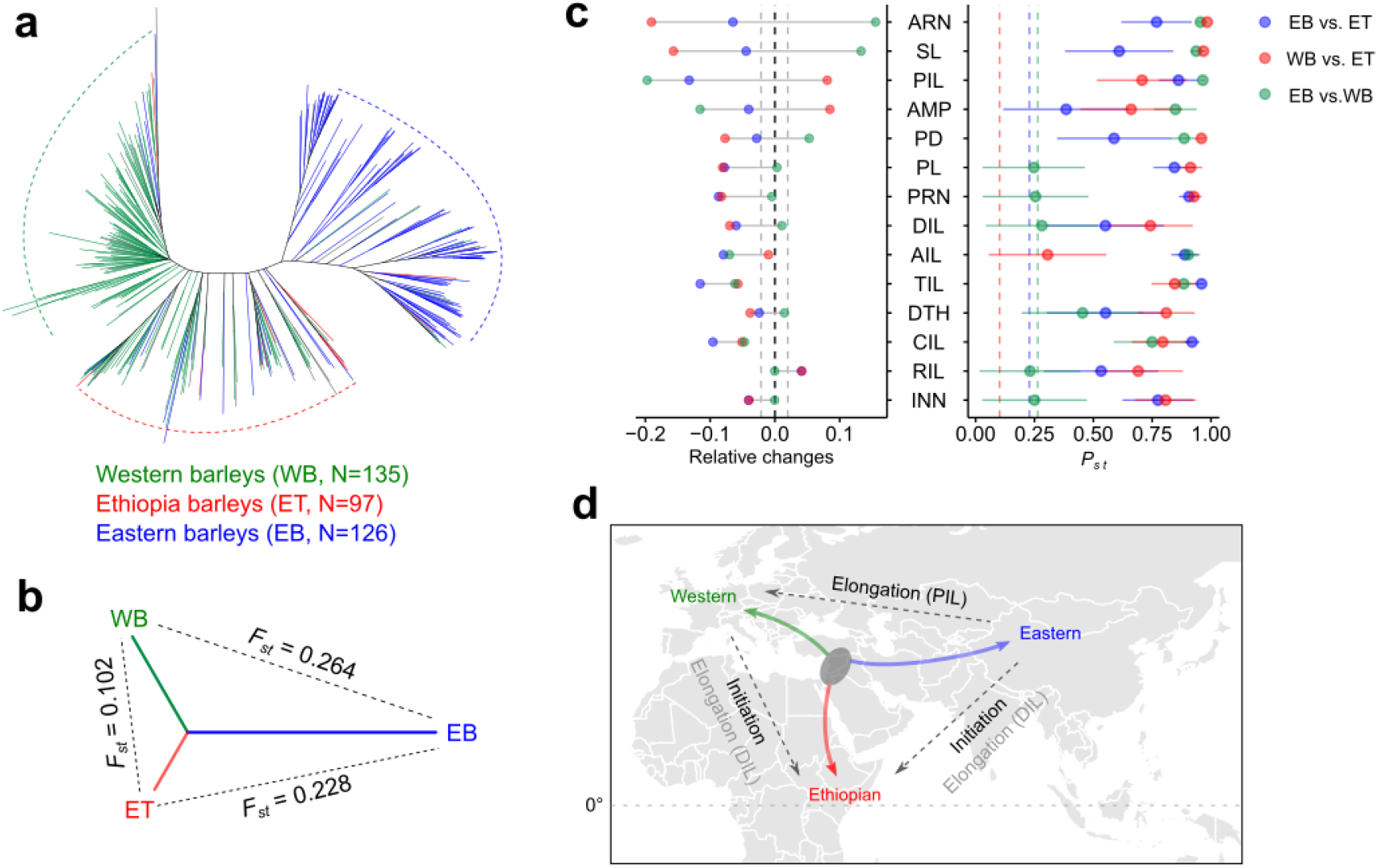
Evolution of phytomer initiation and elongation **a.** An unrooted neighbor-joining tree clusters the 358 barleys (D6S) into three subpopulations corresponding to Eastern (green), Western (blue) and a mix clade (red, Ethiopian clade). **b.** Pairwise comparisons of the genetic differentiation among the three subpopulations based on Fixation index (*F_ST_*). **c.** Pairwise comparisons of the phenotypic differentiation among the three subpopulations. Left panel shows the relative phenotypic changes. Grey dashed lines mark the significant levels determined from two-tailed Student’s *t*-test at *P*<0.05. Right panel shows the phenotypic differentiation (*P_ST_*) versus genetic differentiation (*F_ST_*). Colored dashed lines indicate the between populations mean *F_st_*. Error bars are 95% confidence intervals (CI_0.95_) for the phenotypic *P_ST_* based on 1,000 permutations. EB, ET and WB: Eastern, Ethiopian and Western barleys. **d.** A simplified schematic diagram summarizing how phytomer initiation and elongation are changed during barley evolution based on the current phenotypic dissections. The grey oval highlights the proximate site of the Fertile Crescent where barley was first domesticated. We highlighted prominent phenotypic changes between subpopulations in black. Direction of the arrows point to the subpopulations with higher phenotypic values. Grey horizontal dashed line indicates the equator.

### An oscillatory pattern of internode elongation

Barley culm internodes elongate successively in an acropetal direction, with the distal one (peduncle) being the shortest during spikelet differentiation stages, but becoming the longest after anthesis^3, 12^. The initial assumption was that without exogenous constraints, internode elongation would follow a linear pattern considering the continuity of indeterminate growth^1^. By measuring culm internode length of ∼1,260 plants from the selected 247 ILs (∼5 replicates per genotype) at the anthesis stage, we found that internode length showed a oscillating decline from distal to proximal ends of the main axis. The overall pattern, however, did not follow a purely linear function, but rather displayed an ‘inverse-S’ curve that could be better explained by a nonlinear cubic function (Fig. 1b and Supplementary Fig. 3a-c). Accordingly, culm internode elongation could be broadly trisected into distal, central and proximal compartments. Further examinations of ∼1,025 plants (∼4 replicates per genotype) from the D6S panel demonstrated an overall conserved oscillatory pattern, but with distinct proximal – distal amplitudes intersecting at the central zone of the culm. The wild barley ILs tended to have longer internodes towards the distal end, and shorter internodes towards the proximal end compared to the D6S counterparts, and this pattern is independent of the total node number.

**Fig. 3:**
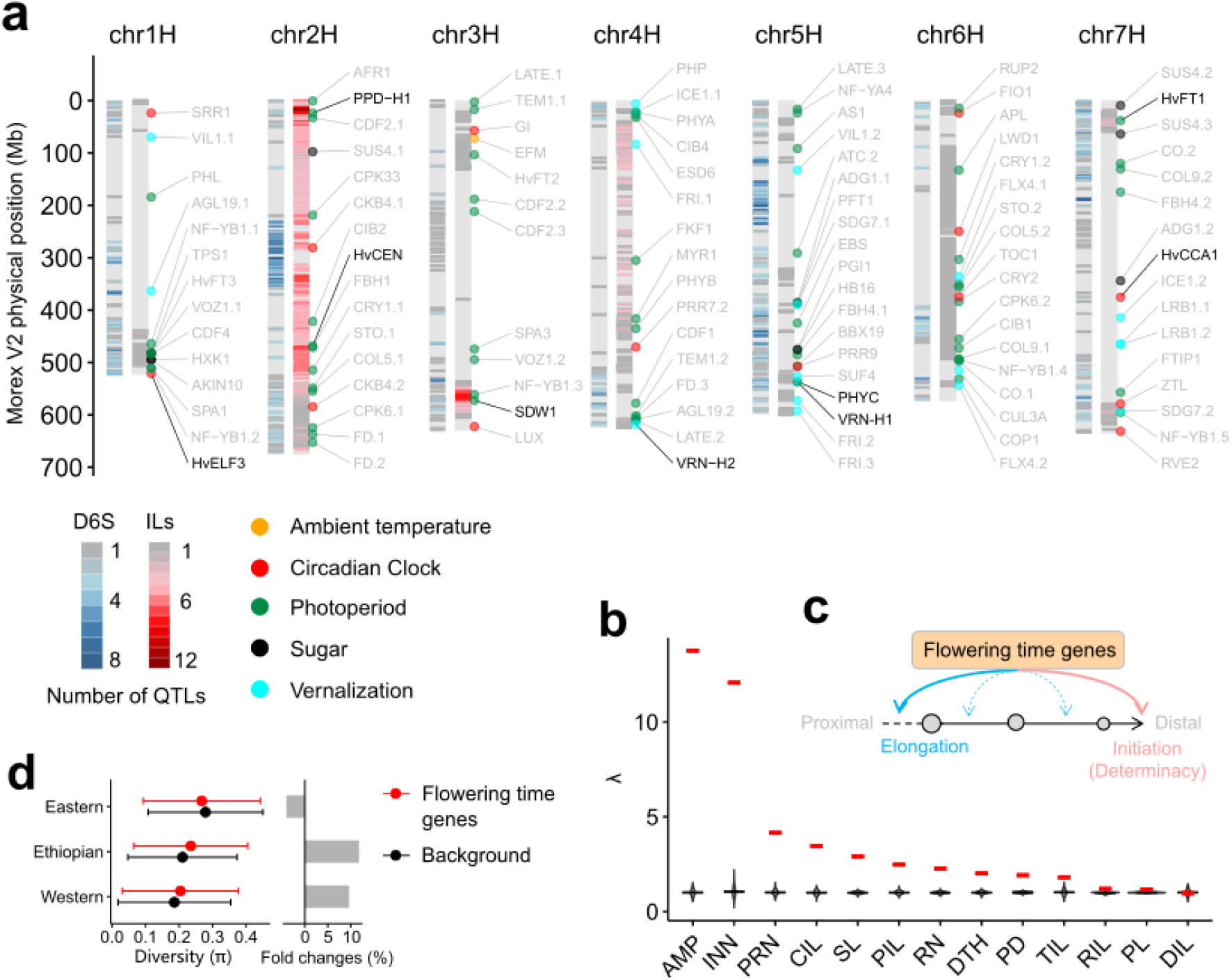
Genetic architecture of phytomer initiation and elongation. **a.** QTL density detected in D6S and ILs for the 14 traits. Barley homologs of *Arabidopsis* flowering time genes from the FLOR-ID database are illustrated along the 7 barley chromosomes. To improve visualization, we only show genes interacting with environmental cues, or circadian clock (112 out of 268), and highlight those that have been experimentally validated in barley with black. **b.** SNP effects of flowering time genes (268 in total) compared to the genome-wide random SNPs. Lambda (λ) values of SNPs within 200-kb of the 268 flowering time gens (280,687 in total) are showed in red, violin plots are the distribution of λ values from 1,000 iterations of random SNPs. **c.** Schematic diagram depicting the functional repurposing of flowering time genes in determining internode elongation based on the summary statistics in (**b**). Flowering time genes are converged on node initiation (or meristematic determinacy^23, 24^, indicated in pink curved arrow), and are repurposed to control internode elongation. Note that the unequal contributions of flowering time genes to different internode elongation (e.g., PIL and CIL) may led to a spatial unevenness of internode length, or higher amplitude (AMP), as observed in (**b**). **d.** Genetic diversity (π) of the 268 flowering time genes in different subpopulations.

Interestingly, based on the measurement of 25 individual spikes from the wild barley ILs, we could also observe an oscillatory elongation pattern for the rachis internodes, but with distinct distributive amplitudes compared to the vegetative culms (Fig. 1c). Collectively, our in-depth phenotypic observations uncovered a previously unrecognized oscillatory pattern for internode elongation, and suggest that culm growth is compartmentalized.

### Phytomer initiation and elongation are highly related

We next examined the relationship between phytomer initiation and elongation. The underlying assumption was that morphogenesis (e.g., the establishment of a phytomer) is composed of hierarchical and sequential developmental cascades, wherein functions of initiation genes can influence the later differentiation and elongation^29–31^. In the barley inflorescence, spikelet (rachis node) initiation and its subsequent growth are molecularly decoupled due to pre–anthesis tip degeneration^23, 32^, we therefore used PRN as a quantitative readout of reproductive node initiation. We estimated the oscillation amplitude (AMP) for culm internodes by considering the deviation of the observed culm length from the expected linear pattern (Methods and Supplementary Fig. 3d,e). Moreover, to better account for differences in node number among accessions, we harmonized each accession by trisecting its culm while applying a moving average strategy to estimate the average lengths of distal (DIL), central (CIL) and proximal (PIL) internodes (Methods and Supplementary Fig. 4). Other phenotypic details were summarized in Supplementary Fig. 1. In total, 14 traits representing node initiation and internode elongation were used, which showed a high repeatability ranging from 0.65 to 0.97, and a broad phenotypic variation (Supplementary Fig. 5, Supplementary Table 1 and 2).

**Fig. 4:**
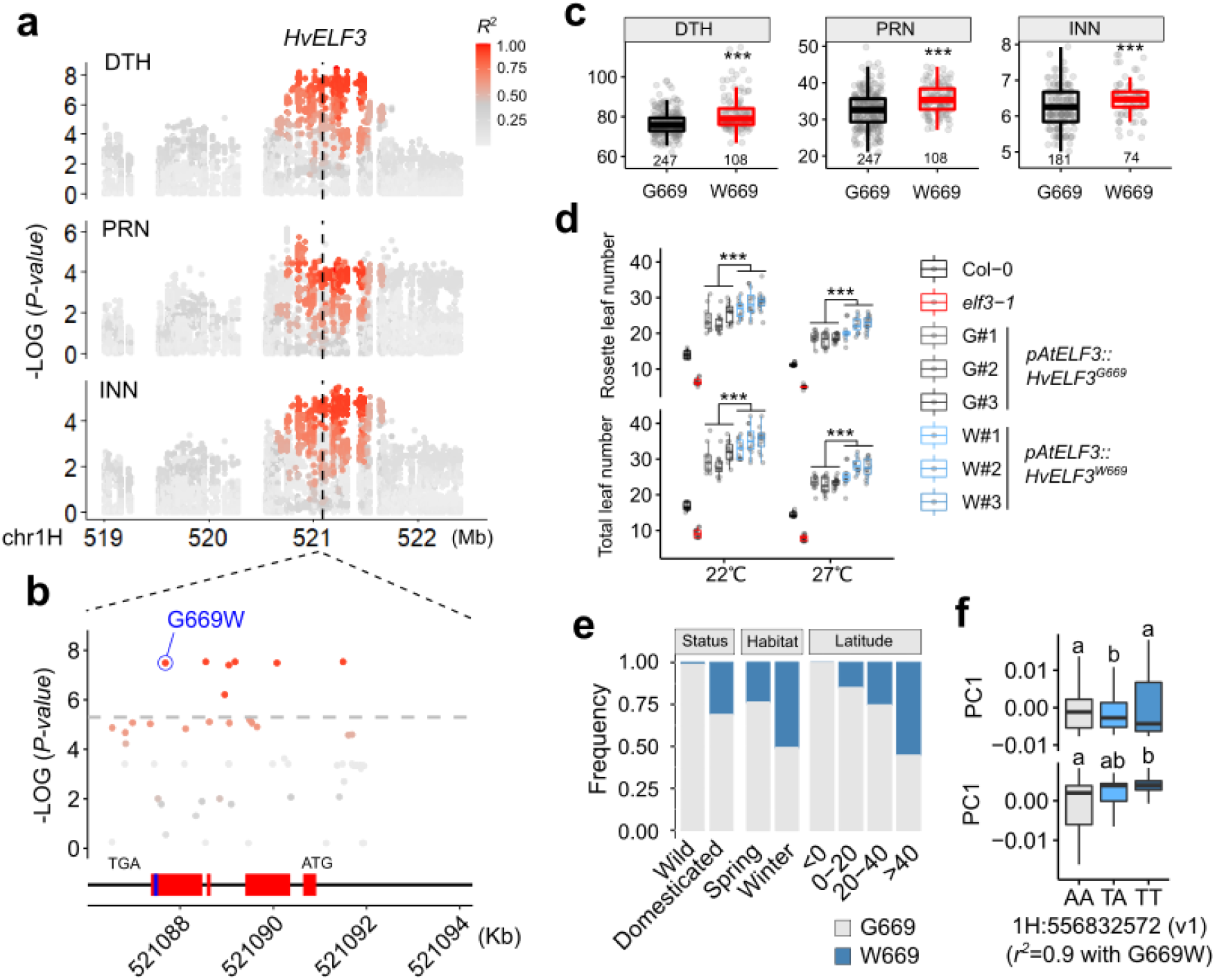
HvELF3 G669W variant contributes to more phytomeric iterations and northward expansion in domesticated barleys. **a – c.** *HvELF3* natural variation is associated with DTH and phytomer initiation (PRN and INN). Local Manhattan plot (**a**) showing the association of the HvELF3 locus on DTH and initiation traits. Relationship (*R*^2^) of the peak SNP with the adjacent SNPs are showed. HvELF3 physical position is indicated with dashed line. A non- synonymous SNP resulting in Glycine (G) to Tryptophan (W) substitution at position 669 of ELF3 protein is highly associated with the traits (**b**), with 669W showing delayed DTH and more phytomeric iterations (**c**). Red boxes indicate exons of *HvELF3*, blue box indicates the putative PrD domain. **d.** Complementation test of HvELF3 variants (669G and 669W) in *Arabidopsis elf3-1* mutant at different temperatures. Three independent events (#1 - #3) for each construct at T_2_ generation are used the phenotypic comparison. **e.** Allelic frequency of HvELF3 G669W in different barley populations or geographical latitudes. We used the 300 re-sequenced barleys, including 100 wild barleys and 200 diverse domesticated barleys^41^, for the comparisons of domestication status (domesticated versus wild) or growth habitat (spring versus winter); we used capital latitudes from the D6S for the latitudinal comparison. **f.** Correlation of HvELF3 G669W variation with eigenvalues of the first two PCs (PC1 and PC2) representing the global domesticated barley diversity in the IPK Genebank collection (n = 19,778)^27^. Note that the direct G669W SNP was not present in the GBS-based SNP matrix (Morex v1), we used the closely linked GBS SNP (1H:556832572, *r*^2^=0.9) as a proxy of HvELF3 G669W. Significant levels in **c** and **d** are determined from two-tailed Student’s *t*-test. \*\*\**P* < 0.001. *n* = 11 or 12 replicates in **d**. Letters above the boxplot in (**f**) represent statistical significance from ANOVA followed by Tukey’s HSD test (*P* < 0.01).

**Fig. 5:**
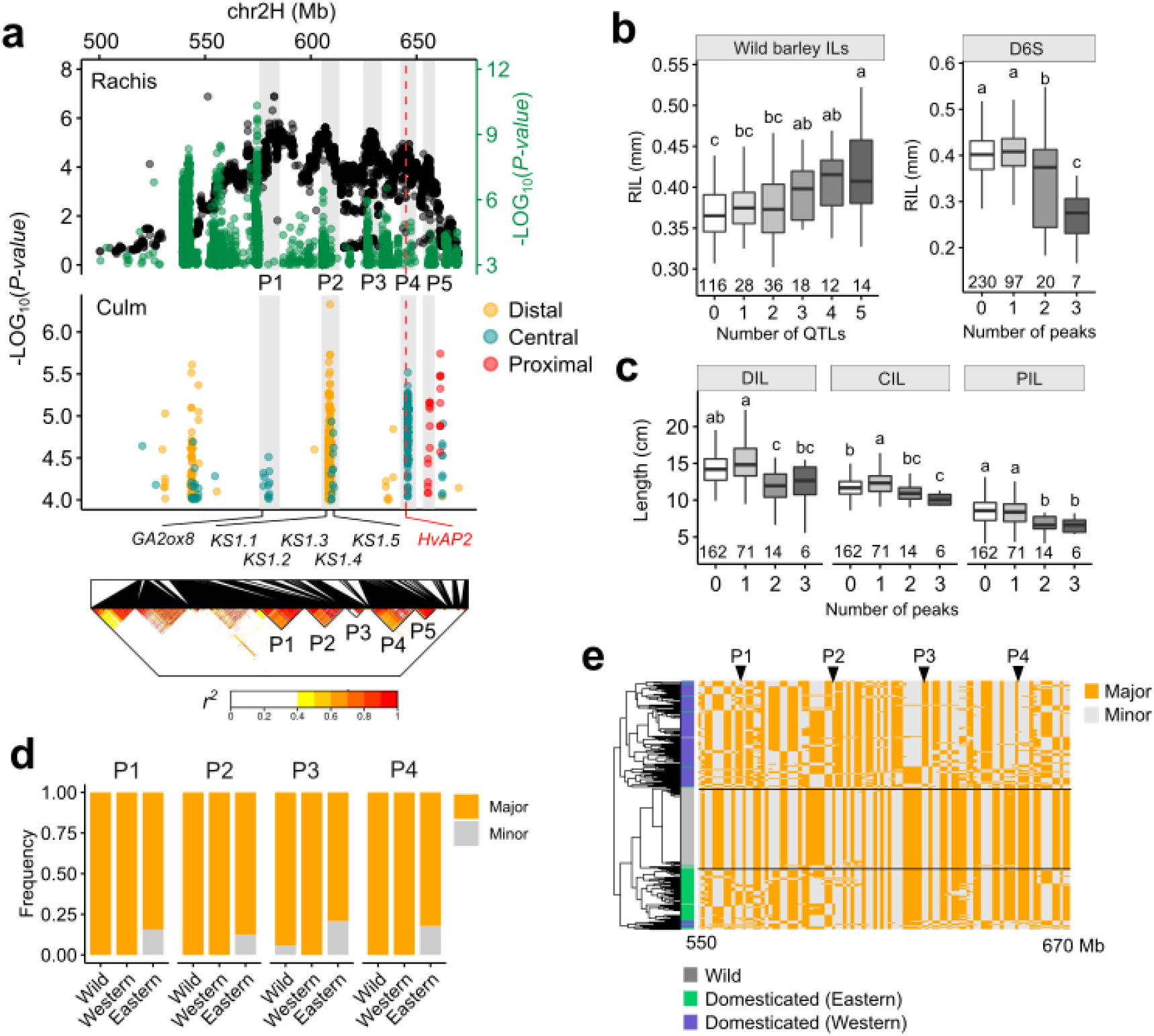
Identification of a super-locus associated with internode length. **a.** Dotplot showing the association signals for internode length. Black and green dots from top panel are association signals detected in the wild barley ILs and the D6S population, respectively. Five adjacent peaks (P1 – P5) supported by both populations are highlighted with grey shadows. *HvAP2* position is marked with red dashed line. Locations of prime candidates involving in GA metabolism are indicated. Bottom panel shows a local linkage disequilibrium (LD) heatmap based on 1,034,167 SNPs (500 – 670-Mb) from the D6S population. **b, c**. Boxplots showing the effects of P1 – P5 on rachis internode length (RIL) (**b**) and culm internodes (**c**). Wild barley alleles additively increase RIL, whereas domesticated alleles additively decrease RIL, as well as culm internodes (DIL, CIL and PIL). Note that no domesticated barleys with a stacking of up to four or five peaks can be identified in the D6S population. Letters above boxplot represent statistical significance from one-way analysis of variance (ANOVA) followed by Tukey–Kramer honestly significant difference (HSD) tests. **d.** Allelic frequencies based on the GWAS peak SNPs from each of the four peaks (P1 – P4) in wild, Eastern and Western barleys. SNP data are extracted from a previous study^41^. **e.** Haplotype diversity within the super-locus in wild, Eastern and Western barley populations. Barley accessions are hierarchically clustered according to 170 binned polymorphic sites from 1,274,896 SNPs (Methods). Locations of P1 – P4 are indicated.

Based on a principal component analysis (PCA) in the ILs, node number and internode length associated traits were largely separated by PC1 (40.2%), suggesting that plants with more nodes tended to have shorter internodes and *vice versa* (Fig. 1d). Indeed, a strong and negative correlation between node number and internode length was observed for both culms (*R*^2^ = 0.34) and spikes (*R^2^*= 0.32). PCA also indicated a high correlation between the growth of vegetative and reproductive phytomers. For example, internode length (*R^2^* = 0.23) and node number (*R^2^* = 0.31) in the culm and the spike were positively correlated (Supplementary Fig. 6a-d). AMP together with several internode length–related traits, further occupied the PC2 axis, which could explain 22.2% of the trait variations. A correlation analysis revealed that only distal internodes (i.e., peduncle and DIL) were significantly correlated with AMP, but not the central or proximal counterparts, indicating that a disproportional elongation of the peduncle might contribute to higher AMP. Indeed, we found that internode lengths were spatially correlated, and that elongation of distal internodes appeared to be largely independent from their proximal counterparts (Supplementary Fig. 6e), revealing a high degree of independency for distal and proximal internode elongation. Finally, peduncle diameter (PD), the outcome of horizontal internode expansion, showed high positive loading on PC1 that was opposite to internode length, indicating an opposing regulation of vertical and horizontal growth.

**Fig. 6:**
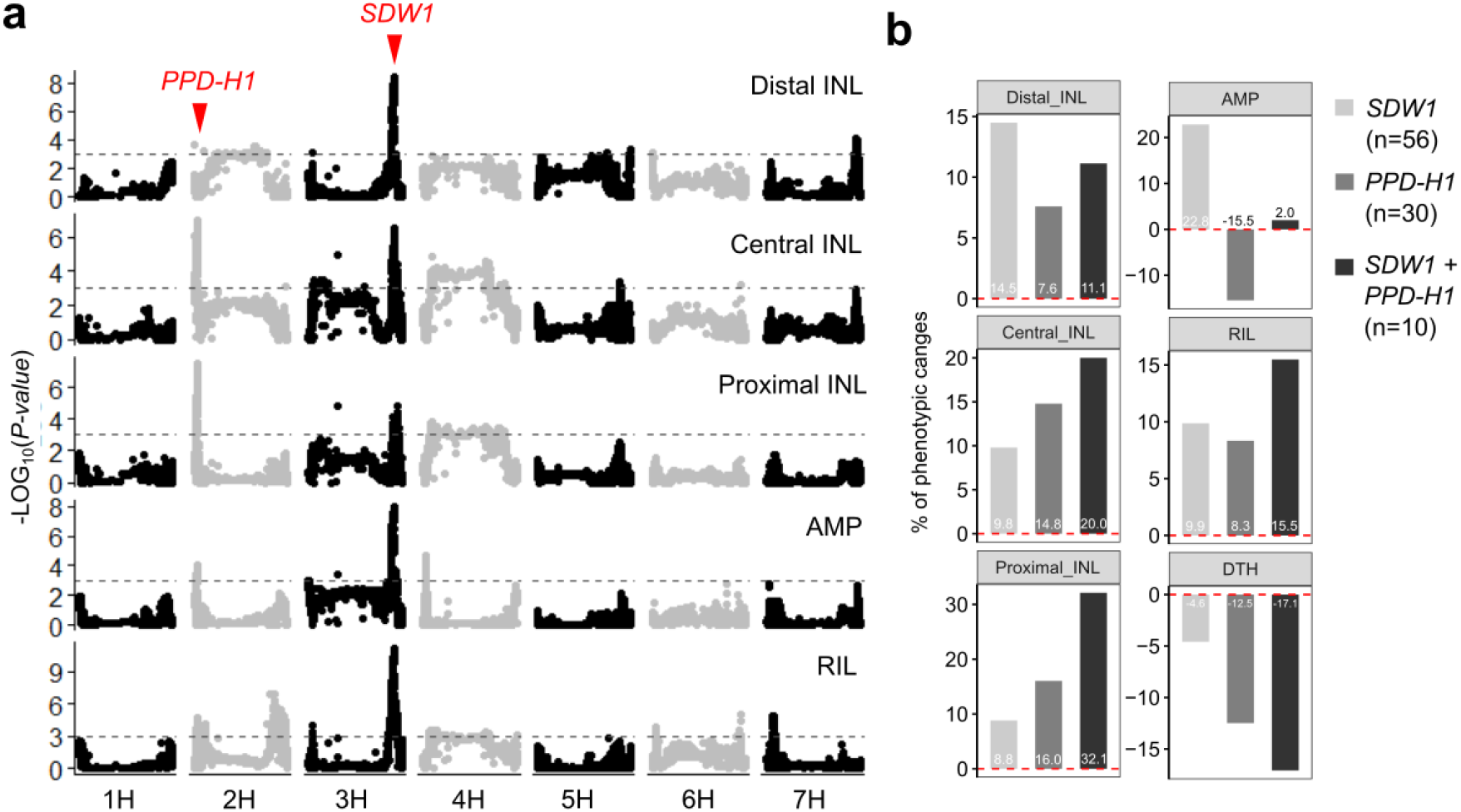
P*P*D*-H1* and *SDW1* loci cumulatively contribute to internode elongation. **a.** Manhattan plots showing the associations for five internode elongation related traits. Locations for *PPD-H1* and *SDW1* are highlighted. Grey dashed lines are genome-wide threshold at *P* = 0.001. **b.** Allelic stacking showing the additive or synergistic effects for *PPD-H1* and *SDW1* on internode elongation, as well as flowering time phenotypes collected previously^26^. Mean phenotypic values from each combination are normalized according to those without wild barley alleles at both loci (n=120).

Because plant height (total culm length, or TIL) is determined by both node number (INN) and internode length, we assessed the relative contribution of INN and each length component (DIL, CIL, PIL) to TIL (Fig. 1e). Using a multiple linear regression, we found that INN, DIL, CIL and PIL together could explain ∼90% (*R^2^* = 92.6% in the D6S and 89.1% in the ILs) of TIL variation. While INN’s contribution to TIL remained similar in both populations, the relative contribution of each length component varied: from the ILs to the D6S, we observed a decline from 41.9% to 21.0% for DIL, and an increase from 3.2% to 21.8% for PIL. Considering the counteracting relationship between node initiation and internode elongation, this result provided evidence that barely can dynamically adjust the growth (both initiation and elongation) of different phytomeric units to stabilize the overall plant height.

### Evolutionary and genetic basis of node initiation and internode elongation

We sequenced (3 – 50 fold coverages) the genomes of the D6S population and obtained 22,405,297 bi-allelic SNPs (minor allele frequency ≥5%), which then clustered the D6S into the three well-defined subpopulations^27^ (hereafter Eastern, Western and Ethiopian barleys) (Methods and Fig. 2a,b). A pairwise comparison of the phenotypes among the three subpopulations demonstrated an overall geographical relevance for the traits assayed (Fig. 2c,d). For example, a latitudinal cline of phytomeric initiation traits (INN and PRN) was observed from the comparisons of Ethiopian barleys with the other two subpopulations, which was likely due to a slight delay in flowering time of Ethiopian barleys under the greenhouse condition. Furthermore, Ethiopian barleys had longer distal internodes (DIL and PL) and higher final spikelet number (FRN) compared with the other two subpopulations. In contrast, the trans-Eurasian comparison (Eastern vs. Western) revealed that, DTH, INN, PRN, as well as DIL and PL, were not significantly different; instead, major difference was observed for PIL, AMP and PD. That said, compared to Western barleys, Eastern barleys had a ‘dwarf and sturdy’ plant architecture due to suppressed proximal internode elongation. Importantly, among the vegetative culm variables, PIL was negatively correlated (*r* = -0.31, *P*=4.24×10^-7^) with spikelet survival, a key reproductive trait that is indicative of grain yielding efficiency^23, 33–35^ (Supplementary Fig. 7). Finally, using a *P_ST_*-*F_ST_* comparison strategy^36^ (Methods), we found that the observed morphological differences among different subpopulations were too large to be explained by random genetic drift (*P_ST_*>>*F_ST_*) (Fig 2c, right), suggesting that selections drove the observed morphological divergences, and thus favored local adaptation.

**Fig. 7:**
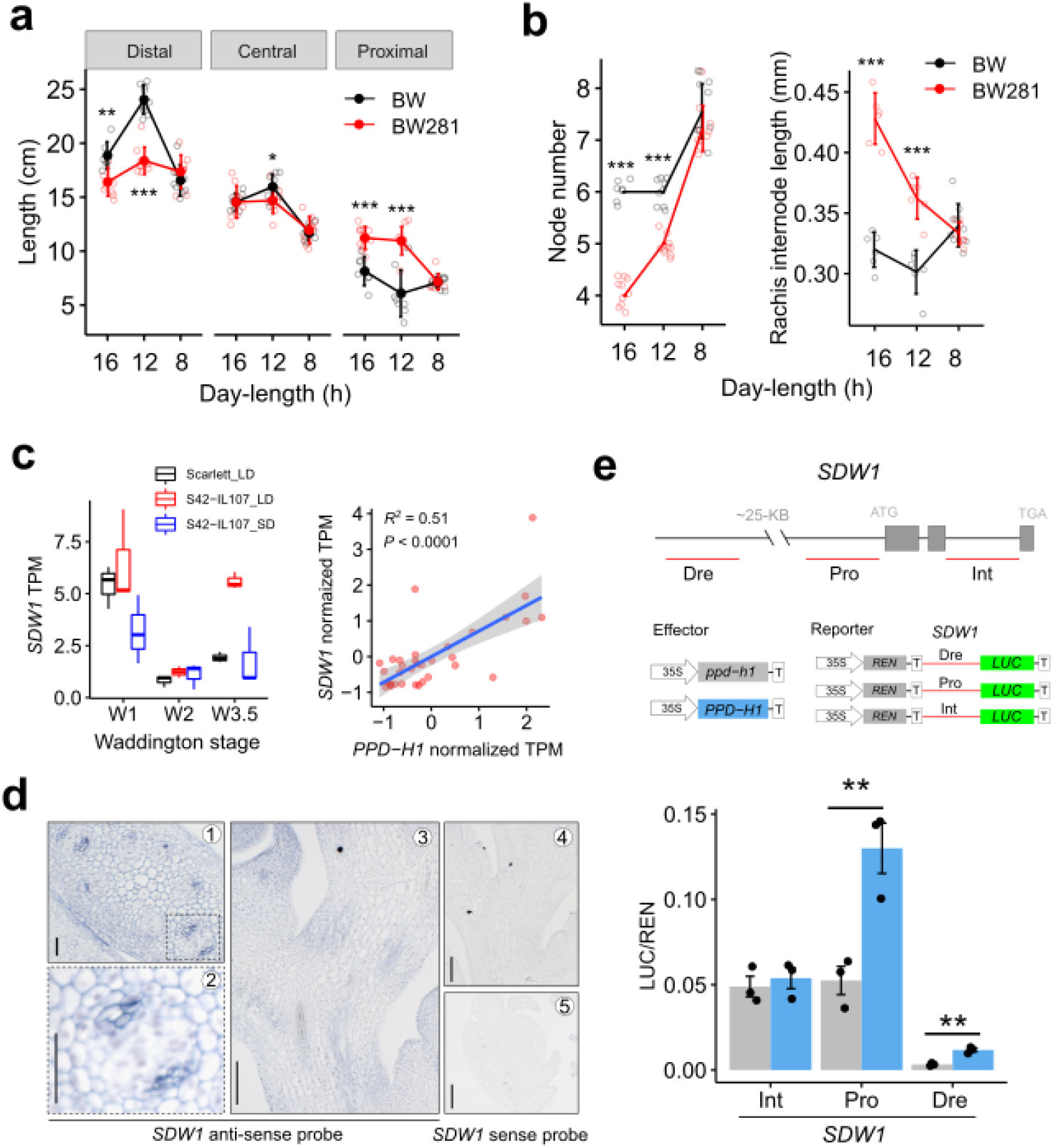
P*P*D*-H1* is molecularly coupled with *SDW1*. **a.** Quantitative comparison of distal, central and proximal internode length under 16, 12 or 8 hours (h) of day-length conditions. **b.** Quantitative comparison of culm node number (left panel) and average rachis internode length (right panel) under 16, 12 or 8 hours (h) of day-length conditions. **c.** PPD-H1 positively regulates *SDW1* gene expression. Boxplot shows the *SDW1* transcripts in developing apexes of a NIL-*PPD-H1* (S42-IL107) and the control (Scarlett) under short day (SD) or long day (LD) conditions. Dotplot shows the co- expression of *SDW1* and *PPD-H1* across diverse tissue-types and developmental stages. RNA-seq data reported previously^49, 50^ are used to estimate the transcripts (TPM) based on Morex annotation v2. **d.** *SDW1* mRNA *in-situ* hybridization in developing rachis during the elongation stage (W5.5) of spike development. Transverse (**d1**, **d2** and **d5**) or longitudinal (**d3** and **d4**) sections hybridized with an anti-sense probe (**d1** – **d3**) or a sense probe (**d4** and **d5**, control) of *SDW1* are shown. Scale bars: 20 μm in **b1** and b2; 50 μm in **d3** – **d5**. **e.** Dual-LUC assay showing the direct activation of *SDW1* by PPD-H1. The diagram on the top depicts the regions used for the assay, including a distal regulatory region (Dre), the 2-kb promoter region (Pro) and the second intron (Int) of *SDW1*. ATAC-seq data reported previously^92^ are used to estimate the accessible chromatin regions of *SDW1*. Relative luciferase activities (LUC/REN) are determined in barley leaf protoplasts co-transfected with different effector and reporter constructs. Data are shown as mean ± s.d. (n = 3 biological replicates). Significant levels in (**a**), (**b**) and (**e**) are determined from two-tailed Student’s *t*-test. \**P* < 0.05; \*\**P* < 0.01; \*\*\**P* < 0.001. *n* = 3 – 9 replicates.

We next sought to understand the genetic basis of these 14 traits by performing a genome-wide association study (GWAS) (Supplementary Table 3 - 5). For the wild barley ILs, we detected 90 QTLs (*P* ≤ 1e^-3^), which explained 5.8 – 52.3% of the phenotypic variances, with wild barley alleles overall reducing node number, but promoting internode elongation in the cv. Barke background (Supplementary Fig. 8). For the D6S panel, we detected 30,363 marker-trait association events (*P* ≤ 1e^-5^), and further clumped them into 468 chromosomal regions ranging from ∼10-kb to ∼10- Mb with a median of ∼2.6-Mb encompassing 2,560 high-confidence genes. Plotting the QTLs from both populations (D6S and ILs) onto the barley chromosomes showed that, their genomic distributions appeared to have high degree of proximity (or overlap), which were preferentially distributed towards the distal ends of each chromosome (Fig. 3a). GO enrichment analysis of the 2,560 genes identified in the D6S indicated that they were mainly involved in biological processes such as photoperiodism, response to cold, hypersensitive response, and GA homeostasis that could be modulated by flowering time genes (Supplementary Fig. 9, Supplementary Table 6). To interrogate this, we conducted a gene-based analysis by focusing on a list of putative flowering time genes according to studies with *Arabidopsis* (268 in total, including several well-known genes highlighted in Fig. 3a, Methods). Based on a lambda (λ) analysis^37^, we found that SNPs located within 200- kb of these genes tended to have higher level of significance for the two initiation traits INN and PRN compared to genome-wide random SNPs (Fig. 3b), confirming the converging roles of flowering time genes in meristematic determinacy and phytomeric iterations^23, 24^. Interestingly, DTH *per se* was not the most significantly associated trait by the variations of flowering time genes; instead, several elongation-related traits including AMP, CIL and PIL, tended to have greater reductions in the *p*- values (Fig. 3b). Because PIL was insignificantly correlated with DTH (Supplementary Fig. 7b, right), these results suggested that flowering time genes could have broader biological relevance for phenotypic variations in internode elongation (e.g., PIL and CIL), which we propose as a functional repurposing (or adopted for a different purpose other than DTH, see below) (Fig. 3c).

**Fig. 8:**
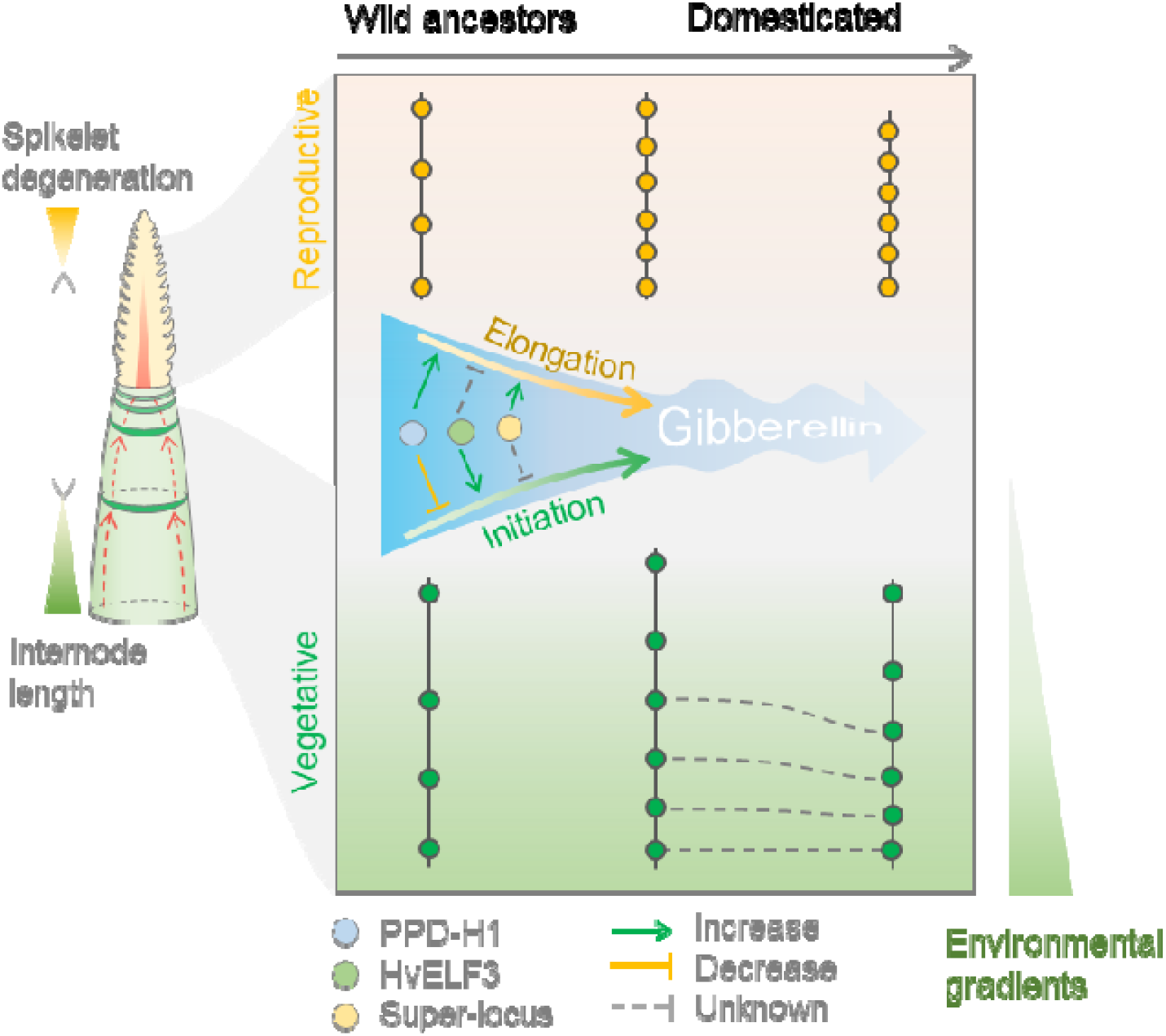
A proposed model depicting how phytomer initiation and elongation are changed under domestication or crop adaptation. During the spikelet initiation/differentiation phase, proximal internode length is positively associated with tip spikelet degeneration, presumably due to extra inputs for elongations or transports (indicated with red dashed arrows). Wild barleys tend to have longer proximal internodes, and early tip degeneration. The transition from this ‘initiated less, elongated more’ of wild ancestor to the ‘initiated more, elongated less’ of domesticated barley can be due to genetic reshufflings of flowering time genes that result in dynamic but overall reduction of GA levels, as demonstrated here three examples (the PPD-H1 – *SDW1* axis, the superlocus, and the HvELF3^18^). The most prominent changes towards the ‘semi-dwarf’ stature of domesticated barleys in Eastern barleys are the shortening of proximal internodes (indicated with dashed lines).

Finally, we found that Ethiopian barleys had a relative higher genetic diversity (π) of flowering time genes than the genome-wide background, whereas Eastern barleys showed the opposite trend (Fig. 3d). This result confirmed the contribution of flowering time gene variations in range-wide geographical adaptation^38^, and suggested a different selective strength of these genes presumably due to contrasting environmental variables (i.e., photoperiod and temperature) across the broad geographical space.

In summary, our genetic analysis identified *a priori* candidates underpinning barley architectural adaptation. Below we detailed three examples to demonstrate an uncoupled relationship for phytomer initiation and elongation: one for node initiation detected in the D6S, one for internode elongation detected in both populations, and the last one for both initiation and elongation detected in the ILs.

### A single amino acid substitution in HvELF3 promotes node initiation during the northward dispersal of barley

One locus on chr1H detected in the D6S panel was found to be highly associated with PRN (*P* = 6.61×10^-7^) and DTH (*P* = 4.03×10^-9^), marginally associated with INN (*P* = 5.98 ×10^-6^), but insignificantly associated with elongation-related traits (Fig. 4a-c). Plants with the minor allele (see below) at this locus were taller than those with the major allele due to more INN (Supplementary Fig. 10a). This locus contained the barley *EARLY FLOWERING3* (*HvELF3*) gene, which encodes a component of the circadian clock acting as a DTH repressor^18, 19, 21^. Previous results indicated that *HvELF3* reduced function mutations accelerated reproductive transition, and facilitated short-season adaptation in 2-rowed spring barleys^19, 21^. However, analysis of the sequence variation at *HvELF3* from the D6S revealed that the GWAS hit was not caused by any of the induced mutations reported previously. Instead, we found a non-synonymous SNP from the top SNPs, resulting in an amino acid substitution from Glycine (G) to Tryptophan (W) at position 669 closed to the C-terminus (here after G669W) (Fig. 4b). This substitution appeared to be located in a putative prion- like domain (PrD) previously found to be essential for thermal responsiveness in *Arabidopsis*, and ELF3 variants lacking the PrD domain showed a constitutively repression of flowering^39^. We found that W669-type had a narrower PrD domain than the G669-type based on a silico prediction^40^ (Supplementary Fig. 10b), and consistently, W669-type plants showed delayed DTH, more PRN and INN compared to those with G669-type (Fig. 4c), suggesting that W669 is a gain-of-function mutation. To test this, we introduced either HvELF3^G669^ or HvELF3^W669^ in the *Arabidopsis elf3-1* mutant^39^. We found that both variants could complement the *elf3-1* mutant phenotypes in terms of leaf number and flowering time at both 22℃ and 27℃ growth conditions (Fig. 4d and Supplementary Fig. 10c). Although no significant Genotype-by-Environment interaction was observed (ANOVA, *P* = 0.24), HvELF3^W6^^69^ transformed plants tended to have better complementation than that of HvELF3^G6^^69^. While other factors may influence the thermal responsiveness, our cross-species complementation data supported G669W as a functional variation.

The G-variant at ELF3 appeared to be restricted to plant species inhabiting at colder climates (Supplementary Fig. 10b). Importantly, we found that the W669 allele was almost absent from wild barleys (1 out of 100)^41^, but had been emerged at higher frequency along with the increase of latitude (northward expansion), in particular, in winter barleys (Fig. 4e), suggesting that W669, which delayed reproductive transition, may facilitate winter survival during northward expanding. Affirming this, wild barley ELF3 alleles (G669-type) accelerate plant development^42, 43^. We extended our analysis to the whole IPK barley Genebank, whose diversity space could be best represented by the first two axes of a PCA corresponding to longitudinal (PC1: Eastern versus Western) and latitudinal (PC2: Ethiopian versus others) gradients, respectively^27^. We found that the G669W variant was highly correlated with the PC2 eigenvalue, but not that of PC1 (Fig. 4f and Supplementary Fig. 10d), supporting an important role of HvELF3 G669W variation for latitudinal adaptation. Collectively, our results indicate that HvELF3 G669W extended the barley phenology (seasonal timing of the lifecycle) to facilitate more phytomeric iterations, and suggest an uncoupled regulation for internode elongation and node initiation/flowering time.

### A super-locus with divergent haplotypes that cumulatively compact plant architecture during the eastward dispersal of barley

We next focused on a large genomic segment encompassing ∼100-Mb mainly associated with internode elongation, such as rachis (in both populations) and culm (in D6S) (Fig. 5a). Multiple physically close peaks (P1 – P5) appeared in this large segment, and notably, peaks for culm DIL, CIL and PIL recapitulated their spatial relationships. For example, shared peaks were detected for DIL – CIL (P2) and CIL – PIL (P5), but not for DIL – PIL. In addition, we also detected both PIL- and CIL- specific peaks, the later was co-localized with *HvAPETALA2* (*HvAP2*), a gene known for controlling both rachis and culm internode length^12, 44^ and was coincided with the P4 region detected in the ILs. A previous study suggested that variations in *HvAP2*’s *microRNA172* (*miR172*) binding site resulted in lower *HvAP2* expression and shorter internode length. However, no sequence variations within the *miR172* region were found from the founder parents of the wild barley ILs, nor did any of the GWAS peaks can be tagged by the *miR172* variations. We speculated that *HvAP2* was unlikely the only causal gene. Indeed, we detected at least eight independent linkage disequilibrium (LD) blocks (*r^2^* ≥ 0.4) within this region in the D6S, including the five shared peaks (P1-P5) (Fig. 5a, bottom). Importantly, we were able to identify diverse allelic combinations in both the wild barley ILs and the D6S, which when more alleles were stacked, showed an additive increase of rachis internode length (RIL) in the wild barley ILs, but an additive decrease of internode length in the D6S (Fig. 5b,c); the additive effect was not observed for initiation traits (Supplementary Fig. 11). This result indicated that multiple independent causal genes for internode elongation were present within the segment, and that domesticated and wild barley alleles likely represented different functional status. Consistently, analysis of the allele frequency based on the GWAS peak SNPs in each of the five blocks showed that the minor alleles were almost exclusively from wild barleys and were preferentially distributed in Eastern barleys (Fig. 5d). We observed a relatively high haplotype diversity across the interval in the domesticated barleys compared to the wild barleys (Fig. 5e), making it difficult to exclude any significant SNPs as tags of other contributing variants. Thus, we postulate that this ∼100-Mb genomic segment is a super-locus that contained multiple independent functional haplotypes (including *HvAP2*) attributing to the mosaic introgression blocks, which cumulatively compacted plant architecture of Eastern domesticated barleys.

Because GA is known to determine internode elongation in a dosage dependent manner, which would fit an additive mode of action observed above^16, 18, 45^, we searched for genes related to GA metabolisms within the interval as *a priori* candidates. We found at least six candidates (Fig. 5a), including one encoding for GA2ox8 homologue in the P1 region, and five for *ent*-kaurene synthases (KSs) participating in the GA biosynthetic pathway^46^ in the P2 region (Supplementary Fig. 12). All of these candidate genes were found to carry either non-synonymous or nonsense mutations specific to Eastern barleys. Similarly, we found Eastern barley-specific mutations in the *HvAP2* gene, suggesting that mutations outside the *miR172* binding site of *HvAP2* could have functional consequence. Taken together, our genetic studies revealed that internode elongation is largely regulated by additive genetic pathways presumably via modulating GA homeostasis; they further showcased multi-functional haplotypes in one super-locus cumulatively contributing to quantitative trait variation, which may have implications for other GWAS – causal gene studies.

### The PPD-H1 – *SDW1* regulatory module coordinates node initiation and internode elongation

Our last example focused on two loci with large effects detected in the wild barley ILs: one for both node initiation and internode elongation, and was coincided with the *PPD-H1* gene on chr2H; the other on chr3H specifically for internode length, and was coincided with *SDW1* encoding a GA20ox2 involved in gibberellin (GA) biosynthesis (Fig. 6a). When using the above-mentioned PCA loadings (PC1 and PC2) as trait variables, these two loci together explained 71.9% of PC1 variation and 54% of PC2 variation (Supplementary Fig. 13a). Both *PPD-H1* and *SDW1* were supposed to be non-functional (or reduced functional) in cv. Barke, but fully functional in all the four wild barleys based on the functional mutations reported previously^20, 47, 48^. Intriguingly, we observed a cumulative shift of the main effects for the two loci on the spatial culm internode elongation (Fig. 6a). For example, wild barley *SDW1* alleles had a major effect on DIL, but *PPD-H1* became the dominant one for PIL variation. Importantly, both loci showed a synergistic epistasis interaction for PIL, and additive effect for RIL or DTH reported previously (Fig. 6b)^26^. The spatial effects on internode elongation for this two loci also applied to the rachises, as functional alleles at both loci promoted longer central-proximal rachis internodes (Supplementary Fig. 13b).

Our genetic analysis also revealed that both *PPD-H1* and *SDW1* loci were significantly associated with AMP, but with opposite effects. Consequently, stacking both loci offset each other’s effect on AMP (Fig. 6b). Because distal and proximal internode elongations were oppositely associated with AMP during flowering (Supplementary Fig. 6e), we hypothesized that proximal internode elongation modulated by PPD-H1 may balance the distal counterparts, resulting in a plant architecture with more evenly spaced nodes and internodes. To further interrogate this, we compared the phenotypes of a *PPD-H1* near isogenic line (BW281; carrying the photoperiod sensitive, functional *PPD-H1* allele) and the wild-type control (Bowman, hereafter BW). Under long-day conditions (16h of light), BW281 overall had shorter plants and spikes mainly due to a severe reduction of phytomer number. However, further measurements for each of the internodes demonstrated that internode length was increased in BW281, in particular, from the proximal ends. Importantly, differences for node initiation and internode elongation among BW281 and BW became insignificant while shortening the day-length (Fig. 7a,b and Supplementary Fig. 13c), suggesting that functional PPD-H1 (BW281) could integrate photoperiod signals to coordinate both node initiation and internode elongation (in particular, the proximal internodes).

The additive or synergistic (i.e., for proximal internodes) genetic relationship between *PPD-H1* and *SDW1* may suggest a molecular interaction of both genes for internode elongation. Indeed, analysis of transcriptomic data generated from developing spikes of cv. Scarlett and S42-IL107 (a Scarlett NIL carrying the photoperiod sensitive *PPD- H1* allele)^49^ revealed that PPD-H1 positively regulated *SDW1* gene expression in a photoperiod-dependent manner (Fig. 7c, left), and likewise, S42-IL107 produced longer rachis internodes than Scarlett (Supplementary Fig. 13b). Moreover, we found that *SDW1* was highly co-expressed with *PPD-H1* in more diverse tissue types comprising a broad range of spike developmental stages^50^ (Fig. 7c and Supplementary Fig. 14a). Interestingly, *SDW1* was found to be more highly expressed in the central sections of developing spikes in both BW and *tip sterile 2.b* (*tst2.b*), a mutant with premature rachis internode elongation^23^. A similar gene expression pattern for *PPD-H1*, however, was only observed in *tst2.b*, which was consistent with the rachis internode elongation patterns along the spike (Supplementary Fig. 14b,c). We previously showed that *PPD-H1* mRNA was detected in the inflorescence vasculature during spike development^23^. Importantly, *SDW1* mRNA *in-situ* signals were also detected in inflorescence vasculatures and rachis internodes (Fig. 7d), which was consistent with its function in rachis elongation. We then tested whether PPD-H1 may regulate *SDW1* expression via interacting with *SDW1*’s regulatory regions. Analysis of chromatin accessibility in *SDW1* revealed several accessible regions that could potentially be the binding targets of upstream transcription factors (Supplementary Fig. 15). In a dual luciferase (LUC) assay, we observed a ∼2 folds increase of the LUC activity from the co-incubations of PPD-H1 with the distal region and the promoter, in respect to those from ppd-H1, albeit the distal region being less transcriptionally active (Fig. 7e), indicating a transcriptional activation of *SDW1* by PPD-H1 during internode elongation.

Altogether, these results demonstrate that the phytomer initiation gene *PPD-H1* is repurposed to regulate internode elongation via the GA pathway acting through the vasculature. This may have broad implications in plant architectural redesign for proximal culm internodes and spike compactness.

## Discussion

In this study, we systematically investigated phytomer initiation and elongation patterns and their genetic underpinnings, and made three fundamental discoveries:

First, compared with rice and maize, inflorescences of domesticated barleys are simplified spike-type inflorescences remarkably resembling their wild progenitor^51, 52^. We found that one key syndrome under domestication is the extension of the apex meristematic phase (indeterminate growth), resulting in more phytomer iterations (Supplementary Fig. 2). This process is coupled with a proportional restriction of the subsequent internode elongation, which is aligned with the general tendency of corps to have more compact stature with more floral organs compared to their wild progenitors^16^.

Second, we showed that phytomer elongation is an oscillatory process and is spatially partitioned into distal, central and proximal compartments. This represents a marked advance from previous work, because we are able to define an underexplored, functional and pillar feature of internode elongation beneath the canopy, whose genetic underpinnings and biological implications are different compared to the distal counterparts (i.e., peduncle). Importantly, this proximal internode growth has undergone a directional selection during barley adaptation. It can be postulated that a similar lengthening rule may apply to other cereal crops, such as wheat and rye, considering the overall conserved phenology of these species^53^.

Third, our genetic analysis pointed to a functional repurposing of flowering time genes for internode elongation. This is because the architecture of all cereal crops, including barley studied here, are segmented into functional phytomeric units. That current floral induction model of mobile signals from leaf to shoot apex is not sufficient to explain the dynamic patterns of different phytomeric elongations, which usually take place after the floral induction. In fact, throughout the lifecycle, many traits such as seed size, that without having direct contacts with the floral induction window, can still be photoperiodically controlled via flowering time gene^54, 55^. The current genetic studies may thus allow a deeper understanding of the dynamic growth strategies in cereal crops.

### Proximal internode as an underexplored functional trait

Architecturally, height increase may confer fitness advantages under natural conditions, such as benefiting pollinations and outcompeting neighbors for light, which is largely achieved through the flowering induced elongation of distal internodes (i.e., peduncle) ^3^. However, under the monoculture farming environment, plant cultivation is usually driven towards the direction of community uniformity and stable yield formation^7^. This would require plants to be able to stabilize their growth in response to the heterogeneous environment at different growth stages. For example, during the pre-anthesis spikelet initiation/differentiation stages, stem elongation usually starts at the proximal internodes instead of the distal counterparts. At this particular developmental window, proximal internodes are spatiotemporally proximate to many unique stress regimes beneath the canopy layer, such as light (discussed below), temperature and moisture. In this context, appropriate switching down of the environmental response machinery, thereby dampening plants’ growth plasticity, may benefit community performance. In particular, shorter proximal internode is associated with higher spikelet survival (Supplementary Fig. 7), and because longer internodes would mean extra inputs^3^, it can be speculated that the shortening of the proximal internodes may enable more resource reallocation to the developing spikes, thereby improving spikelet survival. Another implication for proper proximal internode elongation is its direct relevance to lodging because the bending and breaking of the stems usually takes place near the ground level, and longer internode is considered to be unfavorable for lodging resistance. Since lodging remains a common problem in barley compared to rice and wheat cultivars^8^. Our work may therefore provide a conceptual framework to enhance the genetic gains of proximal internode growth for a sustainable grain yield.

### Light regime as a driving force for dynamic vertical growth

The vertical growth portioning of the internodes immediately suggests light regimes (e.g., red/far-red, R/FR, light ratio) as possible driving forces for selection considering the light gradient at different canopy layers^56^. This is aligned to recent findings demonstrating that increasing canopy light transmission can shorten the proximal – central internodes in both rice and wheat^57, 58^. Indeed, vertical light gradient had already been established at the tillering/elongation stages, and became more evident at the anthesis stage under the greenhouse conditions (Supplementary Fig. 16). In fact, barley PHYTOCHROMEs (PhyA – C) homologs were amongst the candidate list for different internode elongations in the present study (Fig. 3, i.e., PhyA for AMP on 4HS; PhyB for CIL/PIL on 4HL and PhyC for CIL on 5HL). Other photoreceptor genes, including CRYPTOCHROME1 (CRY1), were also found to be within the QTL region for DIL and PL on 2HL. Light perceived by these photoreceptors can induce a series of physiological changes, e.g. the endogenous clock synchronization and photoperiod responses^59^. One scenario is that different internodes may have to synchronize their own clock period to match to the vertical dynamic light gradient, resulting in the oscillatory elongation of internodes. This is aligned to a recent finding that different field microenvironments due to self-shading can lead to an adjustment of the endogenous clock from different leaves, thereby impacting agronomic performance^60^. In this context, the shorter proximal internodes observed in Eastern barleys (many are naked barleys from the Tibetan Plateau) could be due to a local adaptation to the high solar radiation such as seen in the Tibetan Plateau^61^, during which the photoperiodic response machinery has to be switched down^62^. Consistent with this view, our data also suggested that photoperiod insensitivity is coupled with shorter proximal culm internode. The fact that flowering time is insignificantly different between the Eastern and Western barleys under our greenhouse conditions (long-day), may further suggest that the photoperiodic control of internode elongation can be uncoupled from flowering time variation, which is aligned to a recent research in rice demonstrating that photoperiodic response and GA-dependent internode elongation can be uncoupled^63^. In this scenario, the barley flowering time gene *PPD- H1* is repurposed to determine proximal internode elongation via (in part) *SDW1* mediated GA pathway.

### Balancing the phytomer initiation and elongation trade-off to optimize height

Plant height is polygenically controlled in many cereal crop species^64–68^. It is unclear, however, to what extents these height-related QTLs can be explained by the initiation and elongation processes. In the present study using barley as a model, it is shown that initiation (number) and elongation (length) could each explain ∼40% and ∼50% of height variation, respectively. Given the trade-off relationship of initiation and elongation (Fig. 1d), we argue that by measuring the overall adult plant entity as the quantitative readout, it may not be sufficient to recapitulate the whole developmental consequences. Affirming this argument is that the *PPD-H1* gene, which was found to be negatively associated with plant height (TIL) in the present study and in a previous study in wheat^69^, is in fact a positive regulator of internode elongation (i.e., proximal culm or rachis). Our trait-specific case studies of *HvELF3* (initiation) and the super- locus (elongation) may introduce additional levels of complexity for height. While it has been widely accepted that plant height is the outcome of GA level increases, the increasing process can be dynamic. For example, in *Arabidopsis*, GA first acts positively for floral induction, then negatively for inflorescence branching^70^; similarly, the GA signaling protein DELLA negatively controls meristem size independent of height^71^. One legitimate thought is that GA levels need to be repeatedly reset throughout the continuum of plant development, and the timing of this resetting is maintained by the genetic hierarchy of flowering time genes. Thus, genetic reshufflings of beneficial mutations, recurring over thousands of years of crop evolution, have enable a dynamic adjustments of GA levels at different developmental phases or phytomeric units, thereby balancing the initiation and elongation trade-offs (Fig. 8).

In conclusion, our results demonstrate that segmented plants have developed the capacity to heterochronically modify the growth of different functional units (phytomers) during their lifecycles. We propose that plant architecture has to be recognized as dynamic phytomeric units instead of a uniform entity, particularly when studying how plants interact with the heterogeneous environments in the context of crop evolution.

## Materials and Methods

### Plant materials, growth conditions and phenotyping

The 25 wild barley founder parents, together with the 247 wild barley introgression lines, are from a previous research^26^. Wild barley parents for the selected lines are HID003, HID065, HID294, and HID359, respectively. The 358 six-rowed spring barleys were selected from the Federal *Ex-situ* Gene Bank hosted at the Leibniz Institute of Plant Genetics and Crop Plant Research (IPK)^27^, which were described elsewhere^23, 33^, and were supposed to carry functional *PPD-H1* allele based on the SNP22 reported previously^20^. *PPD-H1* sensitive lines (BW281) and S41-IL107 were described previously^72^.

Phenotyping of the wild barley ILs and the D6S population was conducted under controlled greenhouse conditions (photoperiod: 16h / 8h, light / dark; temperature: 20℃ / 16℃, light / dark) at winter season between 2019 and 2022 at the IPK. To phenotype the full set of the D6S panel (358 lines) in a space-limited glasshouse condition, we split the whole panel into two sequential experiments (exp#1, 128 accessions, started at December 2019; exp#2, 256 accessions, started at December 2020), with 25 accessions overlapped between each. We collected only the last two internode length from exp#1, and all culm length data in exp#2. BLUEs and raw phenotypic data of the wild barley ILs and the D6S were given at (Supplementary Table1 and 2). Barley grains were germinated in a 96-well planting tray for 2 weeks, vernalized at 4℃ for four weeks, acclimatized at 15°C for a week, and finally transplanted into 9 cm^2^ square pots until maturity. To estimate the effects of different photoperiods on plant development, barley grains were directly sown in 9 cm^2^ square pots in the growth chambers at three continuous photoperiods (8h, 12h and 16h; temperature: 16℃ / 12℃, light / dark).

We followed an alpha lattice design to control for possible environmental variabilities in the greenhouse, such as different table edges and air conditioner positions. All phenotypic data were collected from the main culm. 4 (D6S) or 5 (wild barley ILs) replicates per genotype were collected. We examined the potential rachis node number (PRN) according to^28^, which was done under a stereo microscope (AxioVision, SE64 Rel. 4.9.1). Final rachis node number (FRN) was counted at anthesis stage. Length-related traits were directly measured with a ruler. Peduncle diameter was measured with a digital caliper. Trait repeatability (*w*^2^) was estimated as 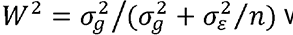 is the genotypic variance estimated by ANOVA, 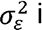 is the residual error variance, and n is the number of replicates per genotype. All phenotypic analyses were done under R (R-3.6.1).

### Estimation of culm internode length variables

The expression form of indeterminacy can be described as a continuous growth process without constraints, which would expect the newly initiated phytomer to be a simple carbon copy (e.g., length, diameter, biomass) of the previous one^1^. Barley shoot apex growth is indeterminate before the transition to the reproductive stage. Thus, the underlying assumption is that the expected length of each culm internode 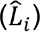 is linearly increased from the proximal (;= 1) to the distal (;= n) ends, which can be expressed as 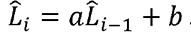. By estimating the degree of deviation (sum of residuals) of the observed length (*L*_*i*_) to the fitted expected length 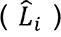(Supplementary Fig. 3c), culm internode length oscillation amplitude (*AMP*) can be estimated as:

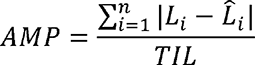

Where *TIL* is the total culm internode length, which was normalized to reduce the effect of plant height (i.e., *AMP* can be independent of height).

Due to a variation of node number among lines, and even within the same line with genetic uniformity, it is not applicable to estimate variations of every internode length. It is also not comparable when considering only the few top/bottom internodes because of the mismatched counterparts caused by node number variation (i.e., second internode is the central one from plants having 4 nodes, but it can be the distal one from those having 8 or 9 nodes). However, based on the elongation patterns in (Fig. 1b and Supplementary Fig. 3), it becomes clear that plants culm elongation follows a trisection rule, regardless of the node number. This trisection rule appears to divide the main culm into three compartments: distal, central and proximal. To facilitate genetic studies, we therefore estimated the average length from the distal (DIL), central (CIL) and proximal (PIL) internodes by using the moving average strategy (Supplementary Fig. 4). To determine the relative attribution of each height component (INN, DIL, CIL and PIL) to total culm length (TIL), we first fit a multiple regression model as described in (Fig. 1e), and then performed the *R^2^* variance decomposition using the Lindeman, Merenda and Gold (lmg) relative importance algorithm implemented in the relaimpo R package^73^.

### Whole-genome resequencing, SNP calling and population structure

High quality genomic DNA was isolated and used for library construction using the Nextera DNA Flex library kit. Library preparation and whole-genome sequencing using the Illumina NovaSeq 6000 device at IPK Gatersleben involved standard protocols from the manufacturer (Illumina, Inc., San Diego, CA, United States). The sequencing adapter sequences and low-quality bases were trimmed using cutadapt (v1.15)^74^. The trimmed reads were then mapped to the Morex reference genome (V2) using minimap2 (v2.20)^75^. Read alignments were sorted by Novosort (v3.06.05) (http://www.novocraft.com/products/novosort/) and then, BCFtools (v1.15.1)^76^ was used to call SNPs, which resulted in 49,526,992 unfiltered SNPs, with 22,405,297 of them having a minor allele frequency of more than 5%.

For population phylogenetic analysis, we first pruned the SNP matrix (22,405,297) using PLINK^77^ (parameters: --indep-pairwise 50 5 0.2), with a window size of 50 SNPs and step size of 5 SNPs, which we obtained 850,398 independent SNPs (r^2^ < 0.2). Phylogenomic tree was constructed based on the distance matrix calculated by PHYLIP 3.68 (https://evolution.genetics.washington.edu/phylip.html), and visualized under iTOL^78^ (https://itol.embl.de/). PCA was performed with PLINK^77^ using the whole SNP data.

### *P_ST_*–*F_ST_* comparison

To investigate whether the degree of the phenotypic differentiation observed between different barley subpopulations could be excluded from the possibility of random genetic drifts, we conducted a *P_ST_*–*F_ST_*comparison^79^. *P_ST_* is a phenotypic analog of *F_ST_*. A *P_ST_* >> *F_ST_* suggests that the observed quantitative trait difference would have exceeded the expectation of the influence due to genetic drift, which is indicative of selection. *P_ST_* is measured as the amount of phenotypic variances explained by genetic group membership: 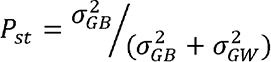 where 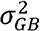 and 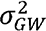 are morphological additive genetic variance components between and within subpopulations, respectively. Because barley is a strictly selfing diploid plant species, which means all polymorphic loci are supposed to be homozygous, there is no need to multiply within-population variance 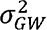 by 2^79^. Calculation of PST was done using the R package pstat v.1.280. We performed 1000 replicates of bootstrap resampling to define the confidence interval of each phenotype. Genome-wide estimates of Fst from the comparisons of different subpopulations were obtained using VCFtools81 by considering the 22,405,297 bi-allelic SNPs (minor allele frequency ≥5%).

### Association mapping

For genome-wide association study (GWAS) in the wild barley introgression lines, we used a simple linear regression model by considering the family information as a covariate, and a significant cutoff as P< 1 x 10-3, which was similar to a previous research82. GWAS in the D6S was done by considering a set of 22,405,297 bi-allelic SNPs. We used a linear mixed model that incorporated pairwise genetic similarities (kinship matrix) as the random effect and additional population structure informed by a principal component analysis (PC1 – 5) as the fixed effect. We ran the GWAS using the software GEMMA83, with a less stringent significant cutoff at P<1 x 10-S.

### Haplotype analysis

LD measurements and visualization for SNPs were done with the R package LDheatmap^84^. To graphically visualize the genotypes within the super-locus (∼100- Mb), we used a sliding window strategy with a window size of 10-Mb and step size of 1-Mb to compress the huge data. In each window, the tested population was forcibly clustered into two groups (assuming all loci are bi-allelic in a diploid species) using the k-means clustering algorithm in R. Major and minor clusters were represented in orange and grey colors in the graphical genotype. Barley accessions were hierarchically clustered according to the resultant genotype of each window (100 windows in total) with the R function hclust. The clustered accessions were colored according to their subpopulation status based on genome-wide SNPs (Supplementary Fig. 12a). A median-joining haplotype network analysis of the 7 prime candidates within the super-locus was constructed and visualized using PopART (v1.7)^85^. For this, SNPs within the genomic regions (from start to stop codons) of each genes from the 100 wild and 200 domesticated barleys reported previously^41^ were used.

### Candidate genes functional enrichment

To summarize the associated candidate genes, significant SNPs for the 14 traits were binned together based on pairwise linkage disequilibrium (LD) decay using the clumping function in PLINK^77^ (parameters: --clump-p1 1x 10-S, --clump-p2 1x 10-4, --clump-r2 0.3, --clump-kb 5000, --clump-allow-overlap). Thus, for every SNP (index SNP) with P < 1 x 10-S, pairwise r2 values were calculated for SNPs within the 5-Mb surrounding the index SNP (±2.5 Mb); SNPs with an r2 2 0.3 and having a P< 1 x10-4 were clumped into bins. Singleton bins without additional SNPs that fell into the criteria were discarded. We recovered 468 bins ranging from ∼10-kb to ∼10-Mb, with a median of ∼2.6-Mb, and encompassing 2,559 high-confidence genes based on Morex reference v2^86^. We used the closest homologs in *Arabidopsis* by considering the highest hit of BLASTP search (e-value < 1e−5) against the TAIR10 dataset. Functional enrichment analysis was done using Metascape (https://metascape.org/)^87^ with default settings. All candidate genes were summarized in Supplementary Table 5.

### Flowering time genes

*Arabidopsis* flowering time genes were extracted from the FLOR-ID database (http://www.phytosystems.ulg.ac.be/florid/) as reported previously^88^, which included 306 experimentally validated genes participating in diverse pathways (latest updated: 2015-09-23). We identified the flowering time gene orthogroups between barley and *Arabidopsis* using OrthoFinder 2.4.1^89^ with default settings. This allowed us to define a set of 268 postulated non-redundant flowering time genes in barley. Major known genes in barley, including *HvFT1* (*VRN-H3*), *HvCEN*, *PPD-H1*, *VRN-H2*, *HvELF3* and *HvCO1* were amongst the list (Supplementary Table 7), suggesting a high degree of conservation on flowering time control. Morex reference has a natural deletion at the *VRN-H2* locus, we postulated the *VRN-H2* physical position using the flanking sequences of the breakpoints, which was estimated to be at ∼618.6Mb in chr4H of Morex v2.

To test for whether variations in flowering time genes were more likely to be associated with phytomer initiation and elongation traits in respect to the genome-wide random subset of SNPs, we used a Lambda (,) analysis^37^:

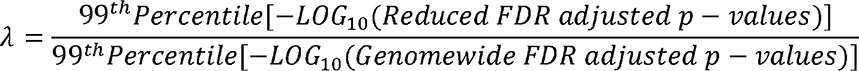

For this, SNPs within 200-kb of the 268 flowering time genes were used (n = 280,687). New False Discovery Rate (FDR)-adjusted *P-values* were calculated using the Benjamini and Hochberg method^90^, and the adjusted top 1% most significant SNPs (99th percentile) were then used to compared with the genome-wide SNPs.

We then repeated the random subset for 1,000 times with the same SNP number, and estimated the , distributions for each trait.

### Cross-species ELF3 complementation

Binary constructs carrying the full length coding sequences of one of the two barley *ELF3* (2298-bp) variants (669G and 669W) and the native *A. thaliana ELF3* promoter (3542-bp) were assembled, and independently transformed into the *elf3-1* mutant^39^ through the floral dipping method. Three independent homozygous transgenic lines for each construct were isolated by phosphinothricin selection.

### Determination of the PPD-H1 – *SDW1* regulatory axis

To examine the regulation of *SDW1* by PPD-H1, we tested the possibility of gene co-expressions using RNA-seq data from previous studies^23, 49, 50^. We used Kallisto software^91^ to estimate the transcript abundances (TPM) of *PPD-H1* (*HORVU.MOREX.r2.2HG0088300.1*) and *SDW1* (*HORVU.MOREX.r2.3HG0256590.1*) using Morex genome annotation V2^86^ as a reference. Putative *cis*-regulatory regions of *SDW1* was estimated with Assay for Transposase-Accessible Chromatin sequencing (ATAC-seq) data reported previously^92^. Data processing, read mapping (to Morex reference V2) and accessible chromatin regions identification were done according to^92^.

We next tested direct regulation of *SDW1* by PPD-H1 using the dual luciferase assay in barley protoplasts. We PCR amplified three potential regulatory regions of *SDW1* from wild-type BW, including the distal region (Dre, ∼1.8-kb, about 50-kb upstream), the promoter region (Pro, ∼2-kb) and the second intron region (Int, ∼1.4-kb), and then cloned them independently into the pGreenII 0800-LUC vector^93^ to generate the *pSDW1^Int/Dre/Pro^*-LUC constructs (reporters). Full-length coding regions (1977-bp) of *PPD-H1* gene from BW281 (functional) or BW (reduced functional) were cloned into the pGreenII 62-SK vector equipped with a 35S promoter (effectors). To isolate barley protoplasts, 7-day-old etiolated seedlings were hand cut into ∼0.5mm pieces with a razor blade, and digested with a freshly prepared enzyme solution (Macerozyme, Cellulase R10, Duchefa) for 6h under darkness with 60 rpm shaking (25℃). Tissues were washed with 50 mL W5 solution [154 mM NaCl, 125 mM CaCl_2_, 5 mM KCl, 2 mM MES (pH=5.7)] 3 times before resuspending with appropriate volume of W5 solution (500 µl per transformation). A 40% PEG-mediated transformation was then applied to deliver the vector combinations into the isolated protoplasts, followed by one more washing step with 880 µl W5 solution, and then resuspended with 1 mL W5. Transformed protoplasts were kept under darkness (25℃) for 16h before the lysis of the cells. Luciferase and renillia luciferase (REN, for normalization) activities were detected with a Dual-Luciferase Reporter Assay System (Promega, E1910) under the GloMax Discover plate reader system from Promega. Primers used in the dual-LUC assays were given in Supplementary Table 8.

### RNA *in situ* hybridization

*SDW1*-specific sequence (402-bp) was PCR-amplified from total cDNA of BW and cloned into the pGEM-T cloning vector. The resulting plasmid was used as a template for preparing sense (negative control) and antisense probes. A fusion primer set containing a 20-bp T7 promoter sequence (5‘- TAATACGACTCACTATAGGG-3’) before the forward primers of sense probes or reversed primer of antisense probes were used. PCR products were then purified and served as templates for in vitro reverse transcription with T7 RNA polymerase. For *in situ* hybridization, spike samples were fixed overnight with FAA (50% ethanol, 5% acetic acid and 3.7% formaldehyde) at 4℃, followed by dehydration with ethanol series (50, 70, 85, 95 and 100%) and then embedded with Paraplast Plus (Kendall, Mansfield, MA). A microtome was used to slice the samples (8 µm thick), which were then mounted onto Superfrost plus slides. Tissue pre-treatment, hybridization, washing and coloration were done as described previously^94^. Primers for amplifying the probe sequences are given in Supplementary Table 8.

### Data availability

For the raw whole genome resequencing reads of the D6S panel (358 lines), 168 will be released in conjunction with an upcoming publication; 47 are from a previous publication^41^; 143 have been submitted to the European Nucleotide Archive database under project id XXX. The unfiltered VCF variant files of the 358 barley lines have been submitted to European Variation Archive database under project id XXX. Genotypic data of the wild barley ILs has been deposited at e!DAL (https://doi.org/10.5447/ipk/2019/20)^95^.

## Acknowledgements

We thank A. Püschel, K. Wolf, M. Schnurbusch and M. Pürschel for their excellent technical support; R. Kamal and N. Shanmugaraj for their dedications in spike dissection and data collection; E. Geyer and his team for their tremendous and tireless support on the greenhouse management; R. Hoffie and I. Otto for assisting the barley protoplast transformation; H. Wang for helps in measuring the LUC/REN activity; S. G. Milner for selecting the barley panel; I. Walde for expert technical assistance in DNA sequencing; Y. Jiang for suggestions on GWAS; A. Fiebig for raw data submission; P.A. Wigge for sharing the *elf3-1* seeds. We are grateful to all members of the Schnurbusch Lab at IPK for fruitful discussions, from which this research concept was stimulated. Research conducted in the laboratory of T.S. received financial support by a European Research Council (ERC) grant entitled ‘LUSH SPIKE’ (ERC-2015-CoG, agreement 681686), the European Fund for Regional Development (EFRE) and the State of Saxony-Anhalt within the ALIVE project (grant ZS/2018/09/94616), the HEISENBERG Program of the German Research Foundation (DFG, grant Nos. SCHN 768/8-1 and SCHN 768/15-1), and the IPK core budget.

## Author contributions

Y.H. conceived the study and performed most of the experiments and data analysis. T.S. supervised the study. Y.H. wrote the first draft of the manuscript and T.S. reviewed & edited the manuscript, with input from all the other authors. A.M. and Y.H. conducted association mapping analysis. R.F.H.G. conducted cross-species ELF3 complementation. G.G. provided intelligence input about internode elongation and crop adaptation. S.Z., V.T., C.T. and Y.H. performed greenhouse phenotyping. S.Z. conducted *in situ* hybridization experiment. C.T. conducted a PCR-based selection of functional *PPD-H1* in the D6S population. G.L. and Y.Z. conducted greenhouse experimental design and phenotypic data analysis. A.B conducted SSD of the D6S population before sequencing. A.H. conducted in-house whole-genome sequencing. M.J. did the SNP calling. N.S. supervised the IPK in-house sequencing platform. M.M. did the selection of the D6S population and WGS analysis. K.P. supervised the construction of the wild barley introgression lines.

---

**Fig. S1.**
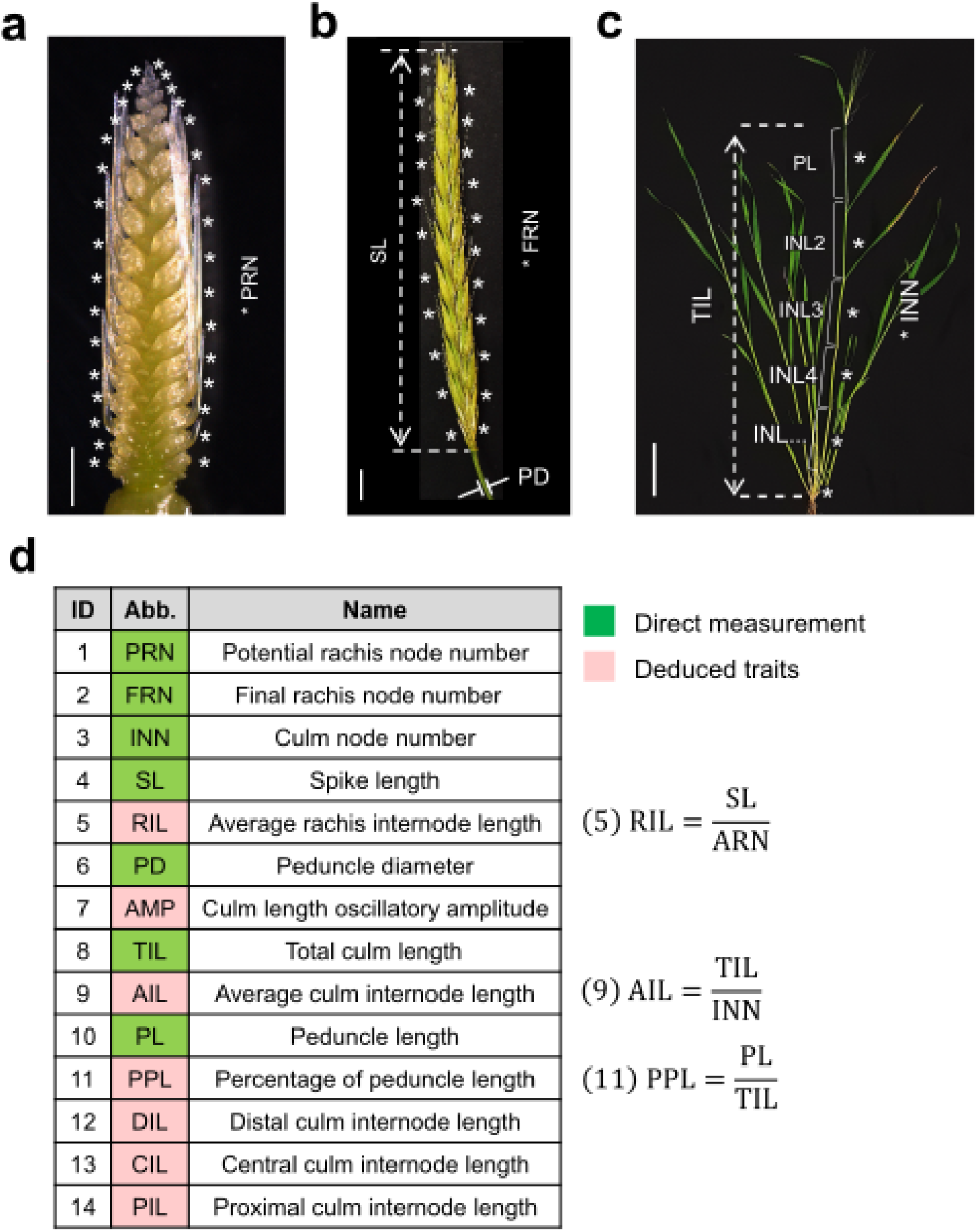
Summary of the phenotypes. **a** – **c**. Representative images showing the phenotyping for node initiation and internode elongation related traits. Reproductive spikes at the maximum yield potential stage (**a**) or anthesis stage (**b**) are used to determine potential rachis node number (PRN) and final rachis node number (FRN), respectively. At anthesis stage, vegetative culms (**c**) are used to collect other phenotypes. Scale bars: 1 mm (**a**), 1 cm (**b**) and 10 cm (**c**). **d**. Trait abbreviations (Abb.) and description of the deduced traits. Other deduced traits (trait id 7, 12 – 14) are summarized in (Supplementary Figs. 3 and 4).

**Fig. S2.**
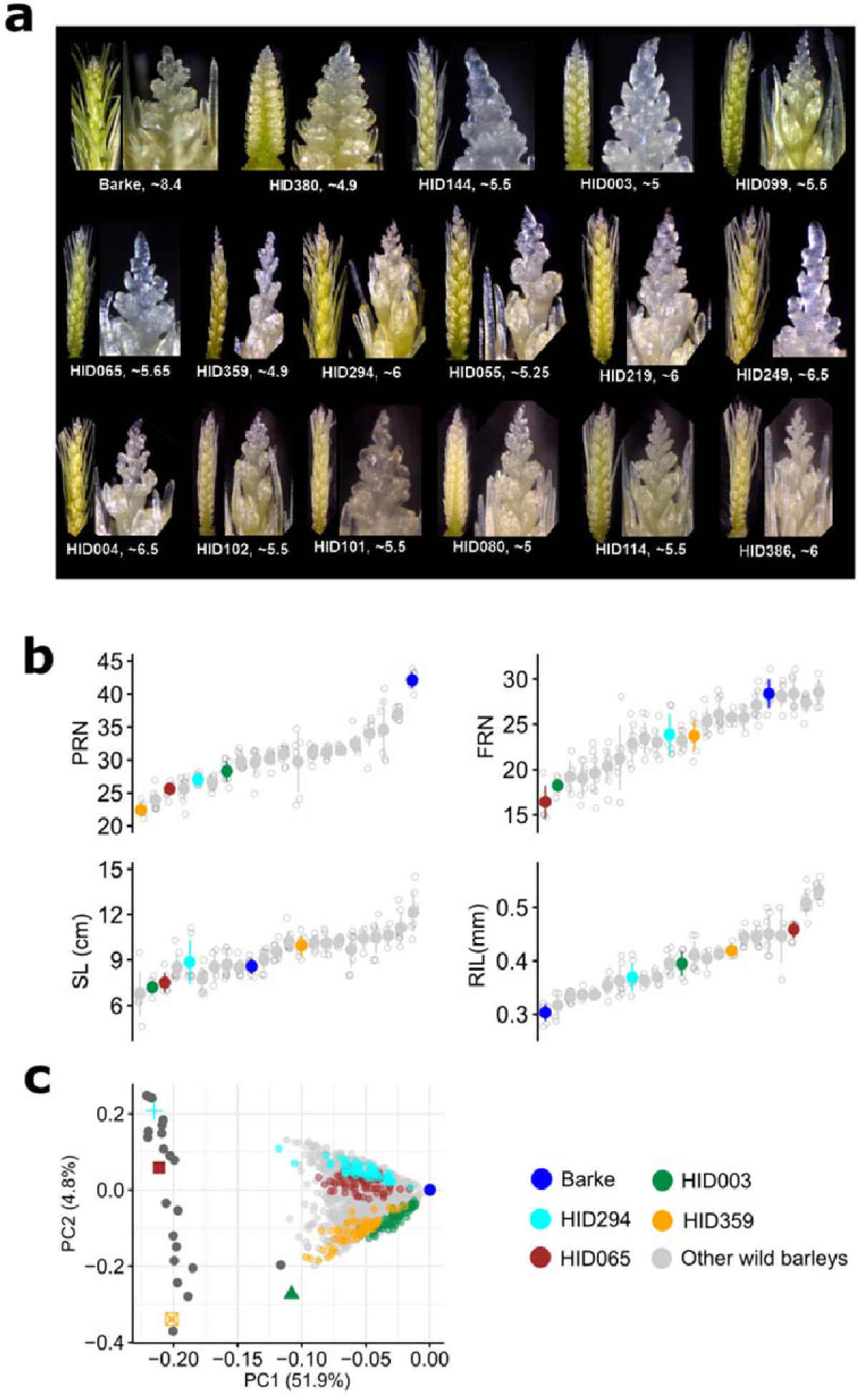
Spike morphology of the HEB-25 founder parents. **a.** Representative spike images highlight the early suppression of apical meristem (inflorescence meristem) activity, and the accelerated tip degeneration from different wild barleys compared with cultivated barley Barke. Numbers below each spike represent spike developmental stage defined by Waddington. **b, c.** Summary of the spike phenotypes from the 25 wild barleys and Barke (**b**). Four selected sub-populations based on the phenotypic and genotypic diversity informed by a PCA plot (**c**) are highlighted with colors.

**Fig. S3.**
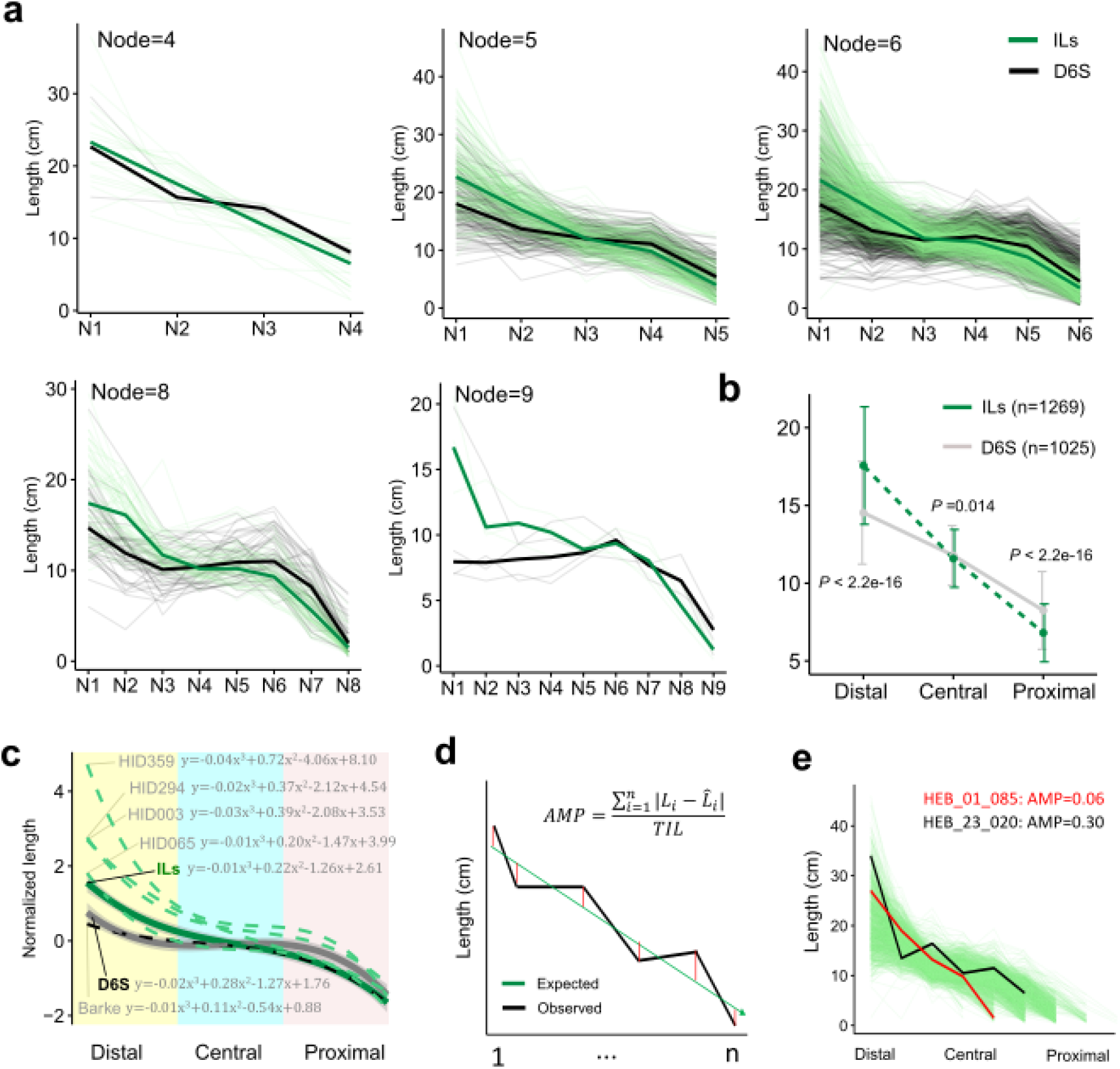
Pattern of internode elongation from vegetative culms. **a.** Pattern of internode elongation from culms with different node number. See also Fig. 1b. **b.** A quartic function modeling the overall patterns of culm internode elongation form the D6S, ILs, as well as the parents. Note that the four wild barley parents overall have both longer distal (DIL) and proximal (PIL) internode length. Compare with wild barley ILs, D6S have shorter DIL, but longer PIL. **c, d**. Estimation of culm oscillatory elongation amplitude (AMP). A graphical depiction of using residuals (Cf, red lines) to estimate AMP. See also Methods. (**d**) illustrates two lines with high (black) or low (red) estimated AMP, in respect to the remaining ILs (green).

**Fig. S4.**
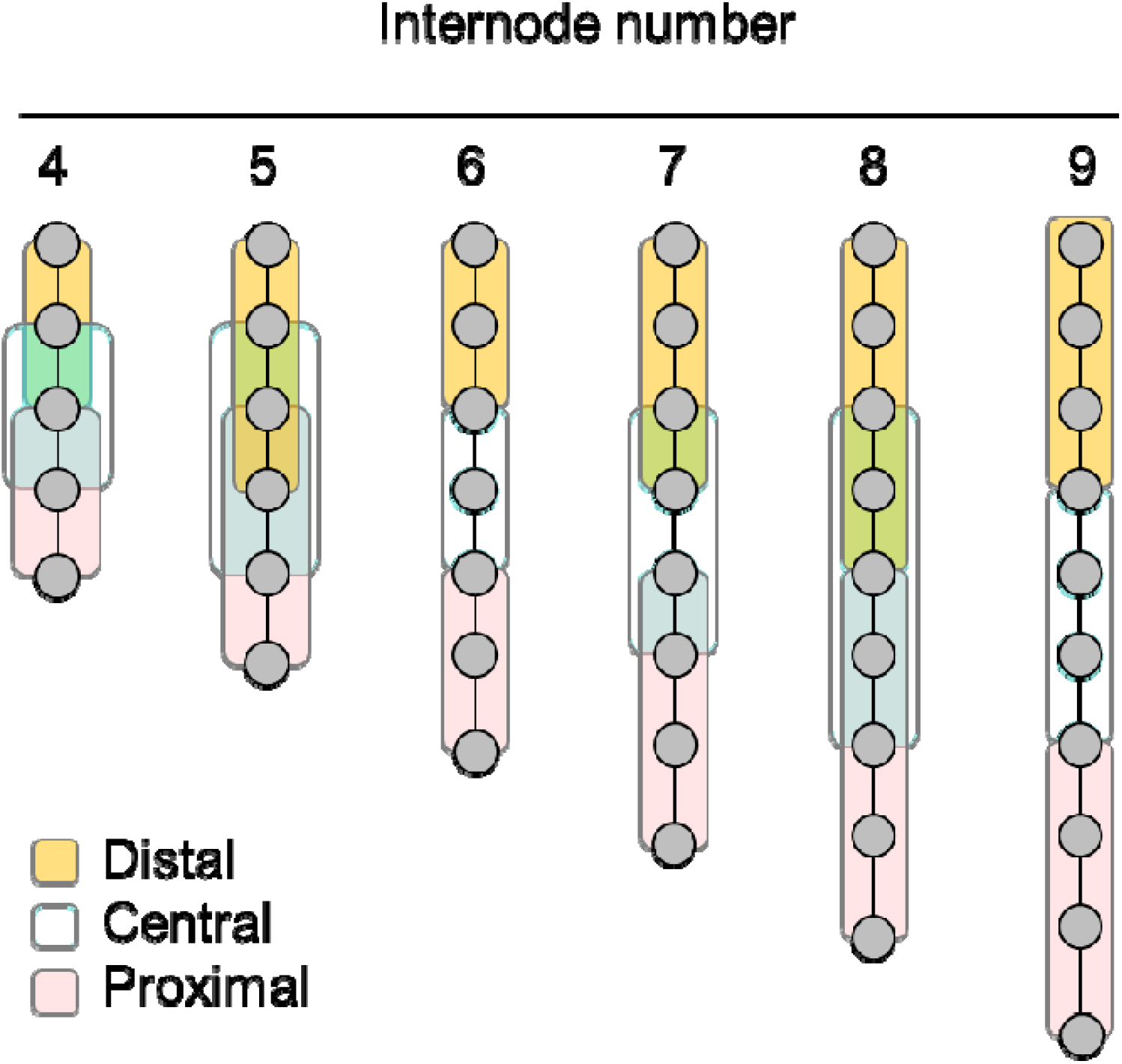
Estimation of distal, central and proximal internode length. A graphical depiction for the estimation of the average distal, central and proximal internode length by using a moving average strategy. Each grey dot represents a node. Lengths of the internodes within each color boxes are averaged to estimate the average length of distal, central and proximal internodes.

**Fig. S5.**
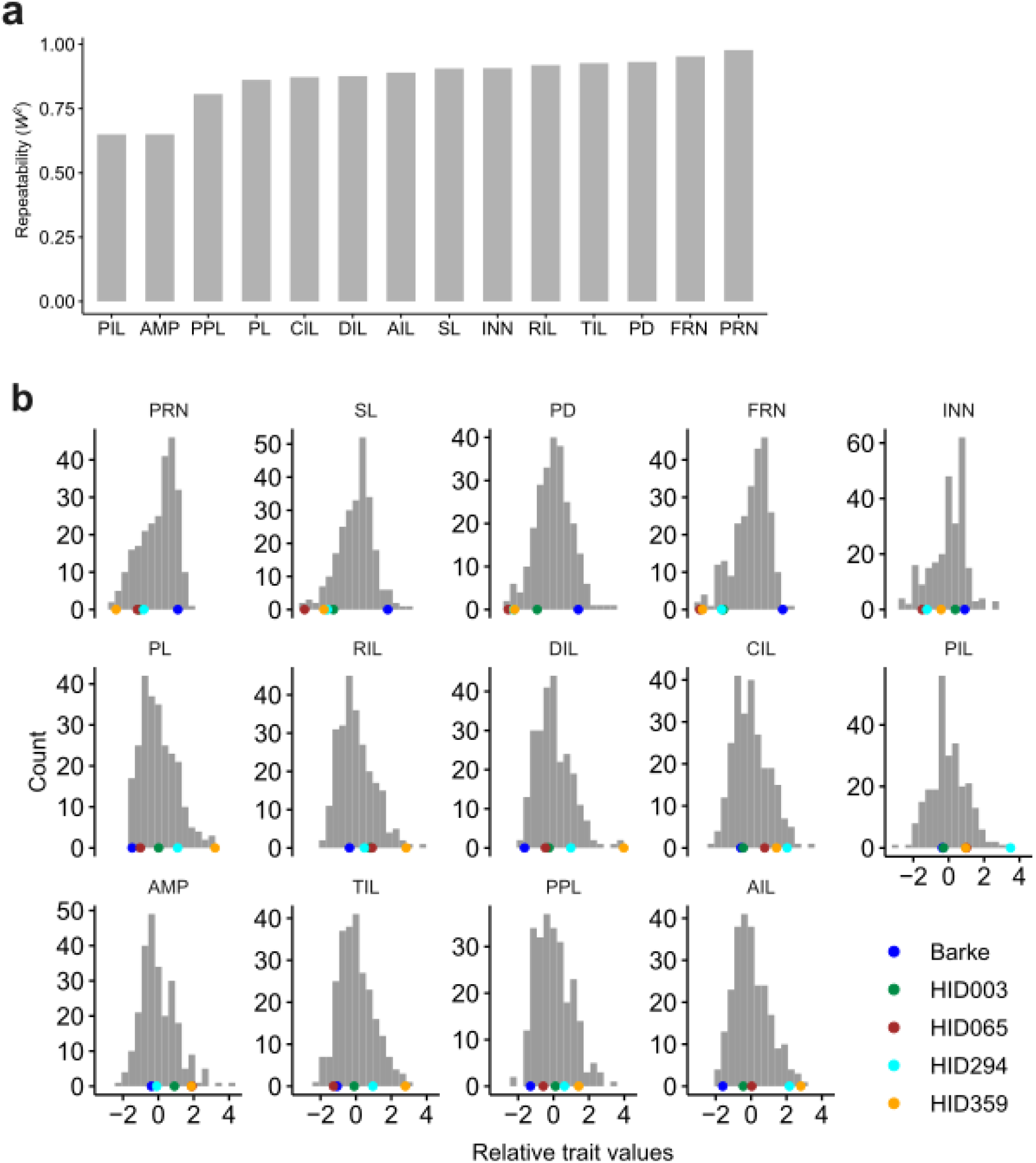
Phenotypic repeatability and variation in the wild barley ILs. **a.** Bar plot showing the estimated repeatability (*W*^2^) for the 14 traits (5 replicates). **b.** Histograms showing the variation of the 14 traits in the wild barley ILs. Parental trait values are highlighted with color dots on the bottom of each histogram. A z-score normalized phenotypic values are shown.

**Fig. S6.**
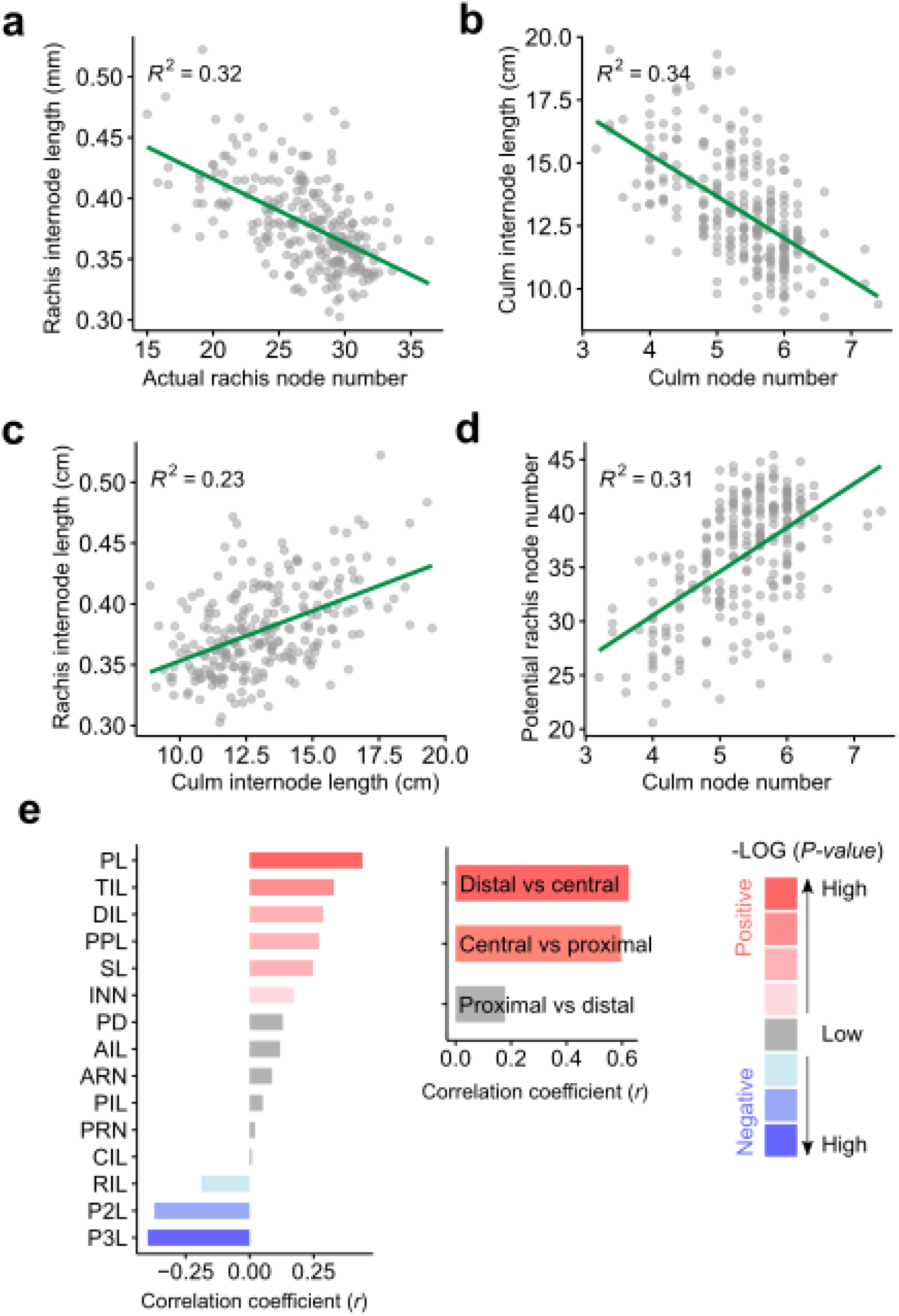
Phenotypic relationship for node initiation and internode elongation. **a – d.** Graphs showing the overall liner relationship from the comparisons of node initiation versus internode elongation (**a**, reproductive; **b** vegetative), or vegetative growth versus reproductive growth (**c**, internode length; **d**, node number). **e.** Relationships of AMP with other phenotypes (left) or among the distal – central – proximal internode length (right) based on the Pearson’s correlation coefficient (*r*). Grey color indicates an insignificant correlation between the traits.

**Fig. S7.**
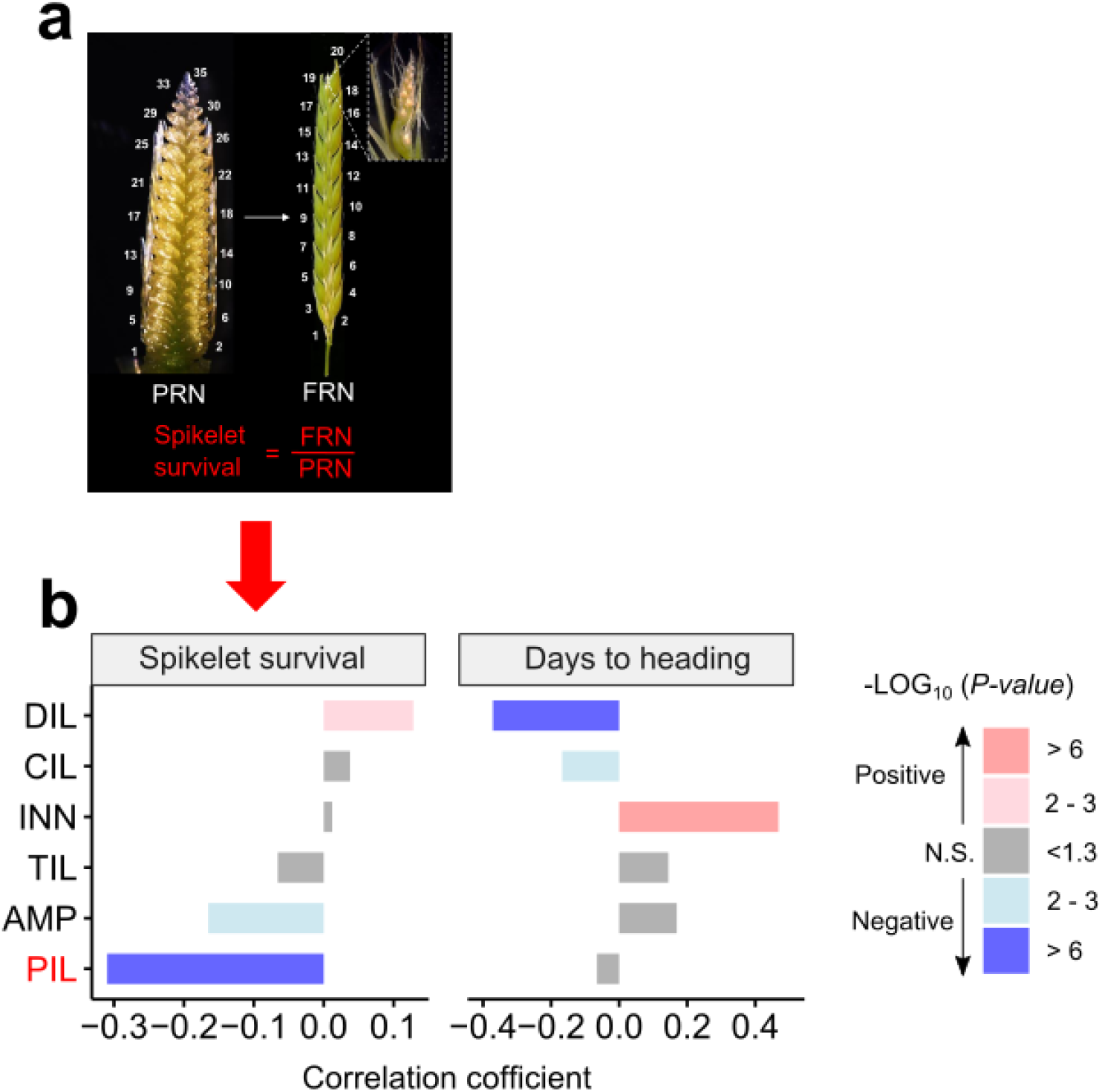
Correlation analysis of vegetative culm variables with reproductive efficiency Left panel depicts the calculation of reproductive efficiency, which is deduced from the fraction of final rachis node number (FRN) by potential rachis node number (PRN). Right panel shows the correlation coefficient of reproductive efficiency with different vegetative culm variables, including distal (DIL), central (CIL) and proximal (PIL) internode length, internode number (INN), total culm length (TIL) and the lengthen amplitude (AMP). Note that among these vegetative culm variables, PIL shows the strongest negative correlation with spikelet survival. Grey color indicates insignificant correlation.

**Fig. S8.**
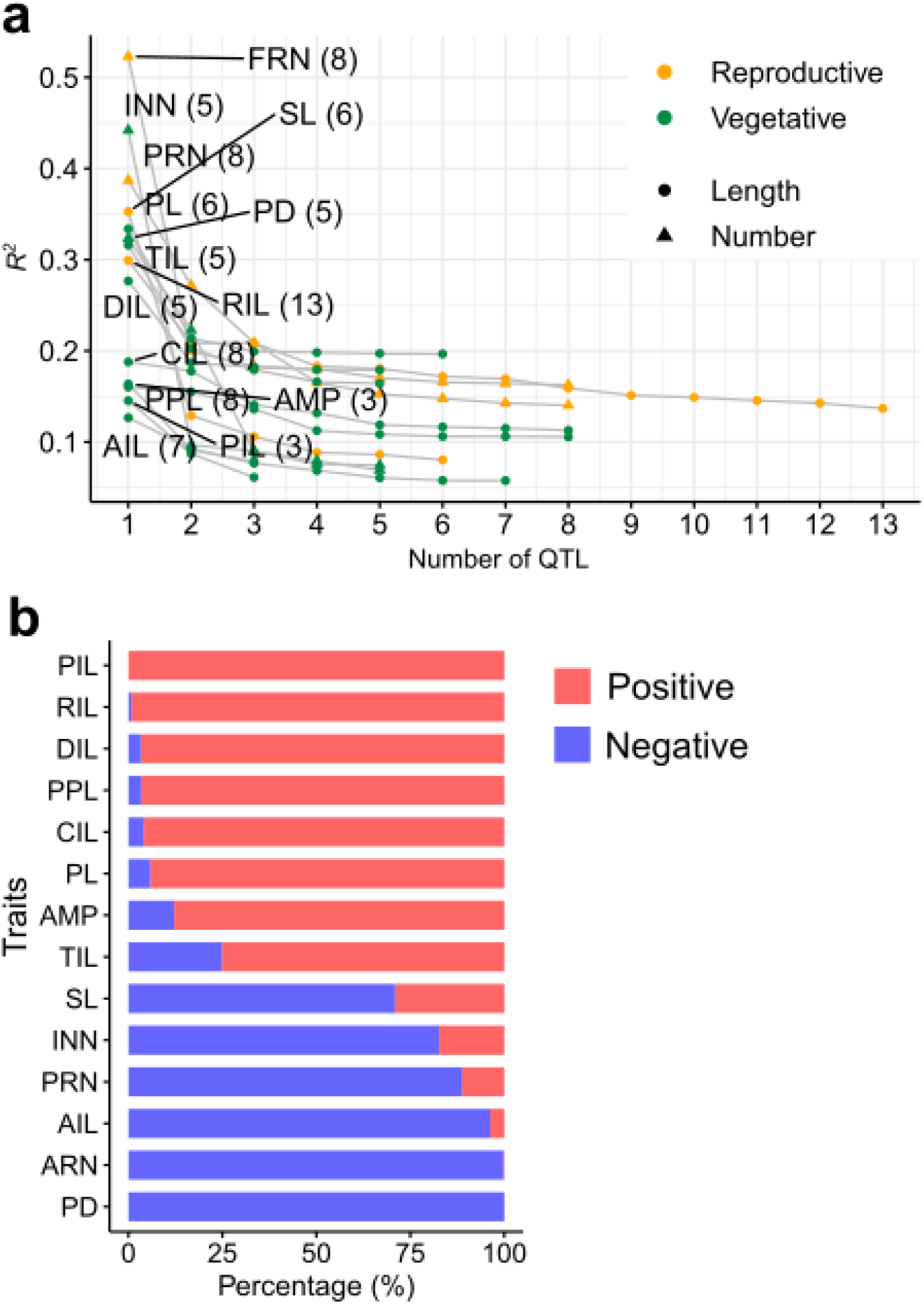
Summary of the quantitative trait loci (QTLs) identified in the ILs. **a.** Distribution of phenotypic variation explained by each QTL (*R*^2^) and the number of detected QTLs. **b.** Stacked bar graph showing the SNP effects on the phenotypes assayed. Positive or negative represent wild barley alleles will positively or negatively affect the traits under Barke background, respectively.

**Fig. S9.**
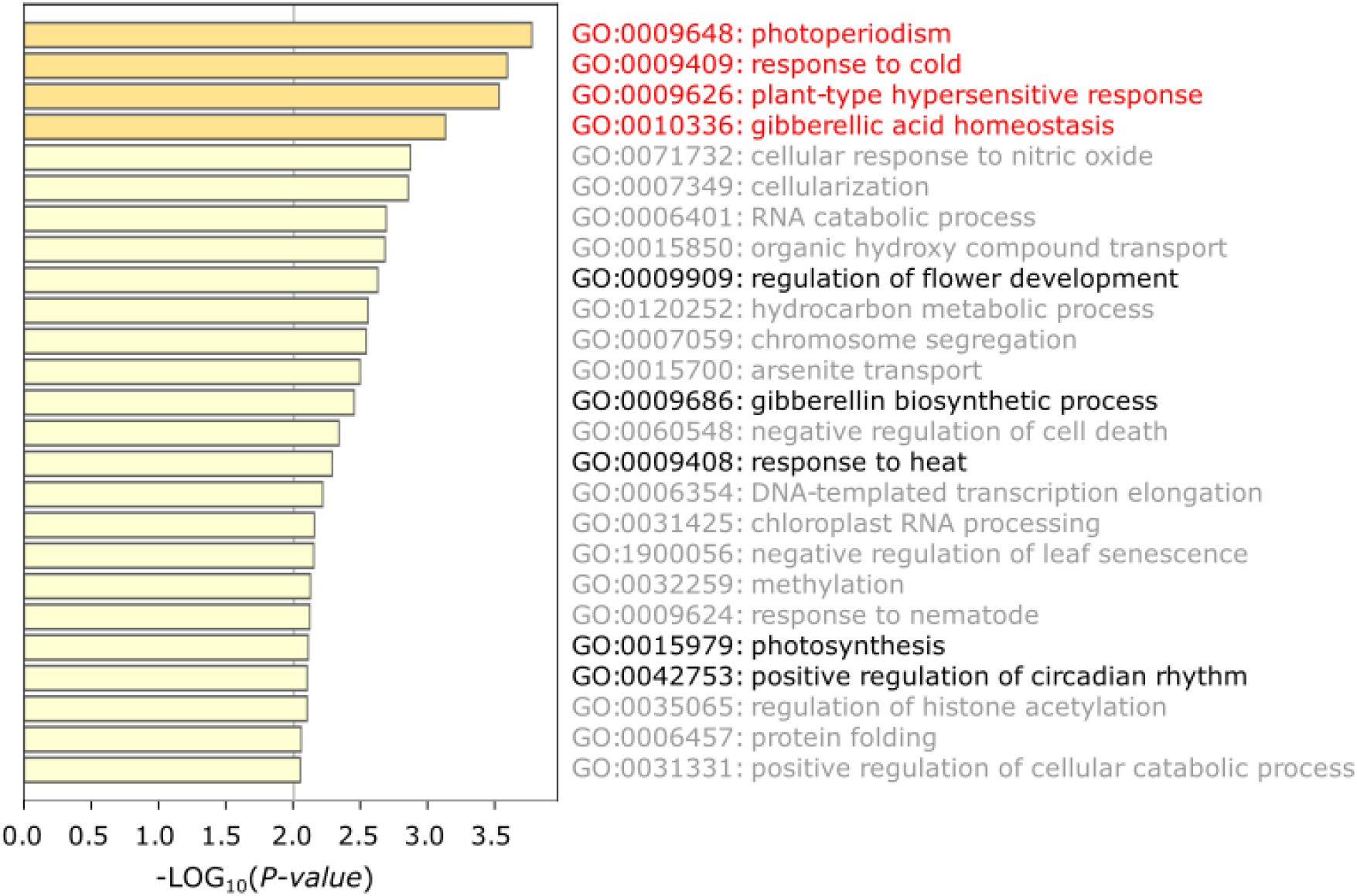
GO enrichment analysis of 2,560 GWAS candidates identified in the D6S The closest barley homologs of *Arabidopsis* genes were first identified through a BLASTP search. Only the best hit of each gene (e-value<1e-05) was used. Functional enrichment analysis was done with Metascape (https://metascape.org). The top 4 enriched terms are highlighted with red color, other less significantly enriched terms potentially related to adaptation are highlighted with black.

**Fig. S10.**
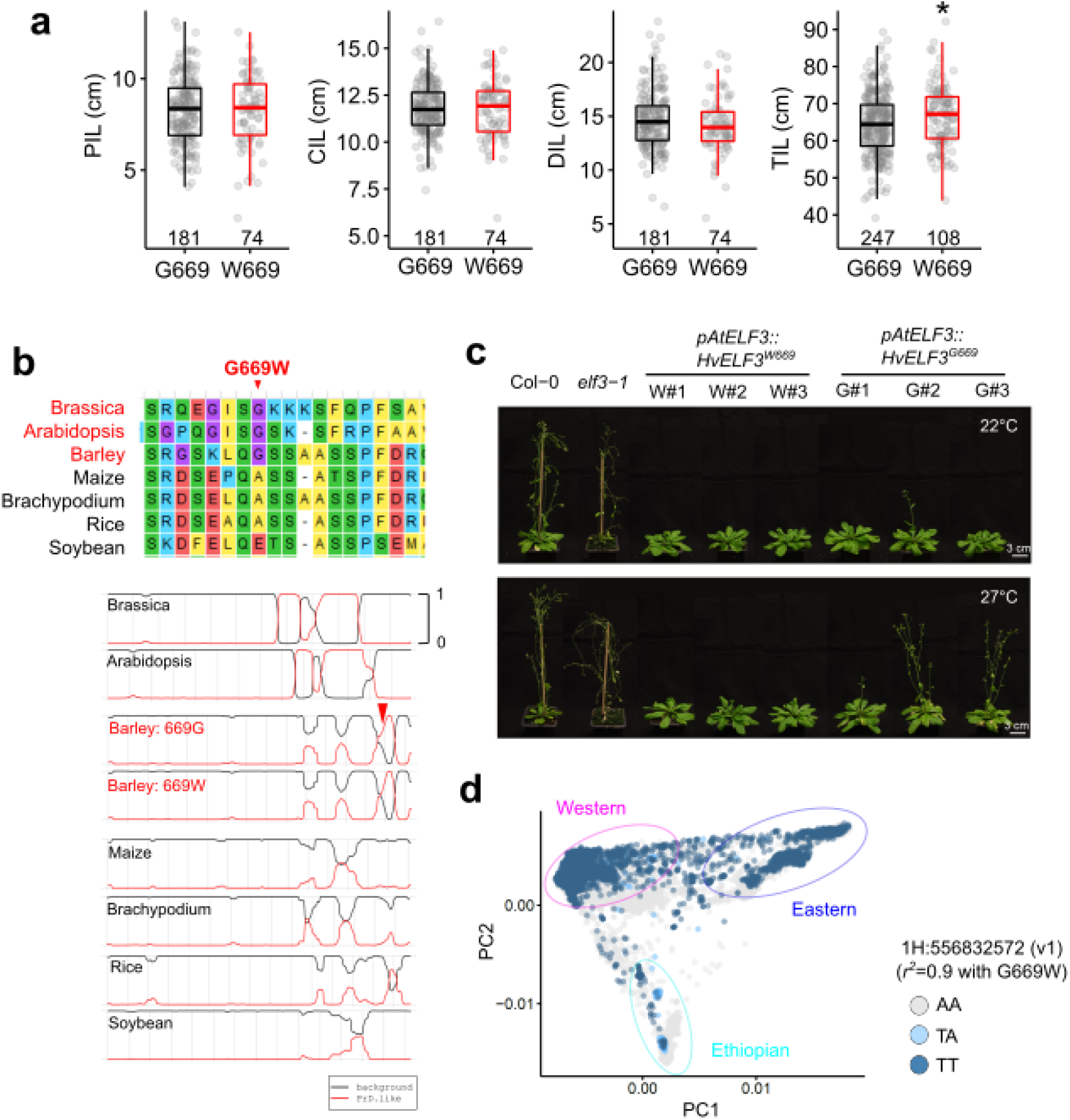
HvELF3 G669W variant and the functional consequence. **a.** Comparisons of internode elongation and total culm length. Note that only total culm length (TIL) was significantly changed (\**P*<0.05) due to the G669W substitution, but not internode elongation traits. **b.** The G669W is predicted to slightly narrow the prion domain (PrD). In silico prediction was done using the Prion-Like Amino Acid Composition (PLAAC) algorithm (http://plaac.wi.mit.edu/). Top panel shows the sequence alignment surrounding the G669W mutation sites in 7 plant species with (red) or without (black) the PrD. **c.** Representative image showing the differential complementation of flowering time for the *Arabidopsis elf3-1* mutant with barley HvELF3^669W^ or HvELF3^669G^ variants. **d.** HvELF3 G669W variation is mainly associated with the PC2 axis that separate Ethiopian barleys from the remaining. See also Fig. 4f.

**Fig. S11.**
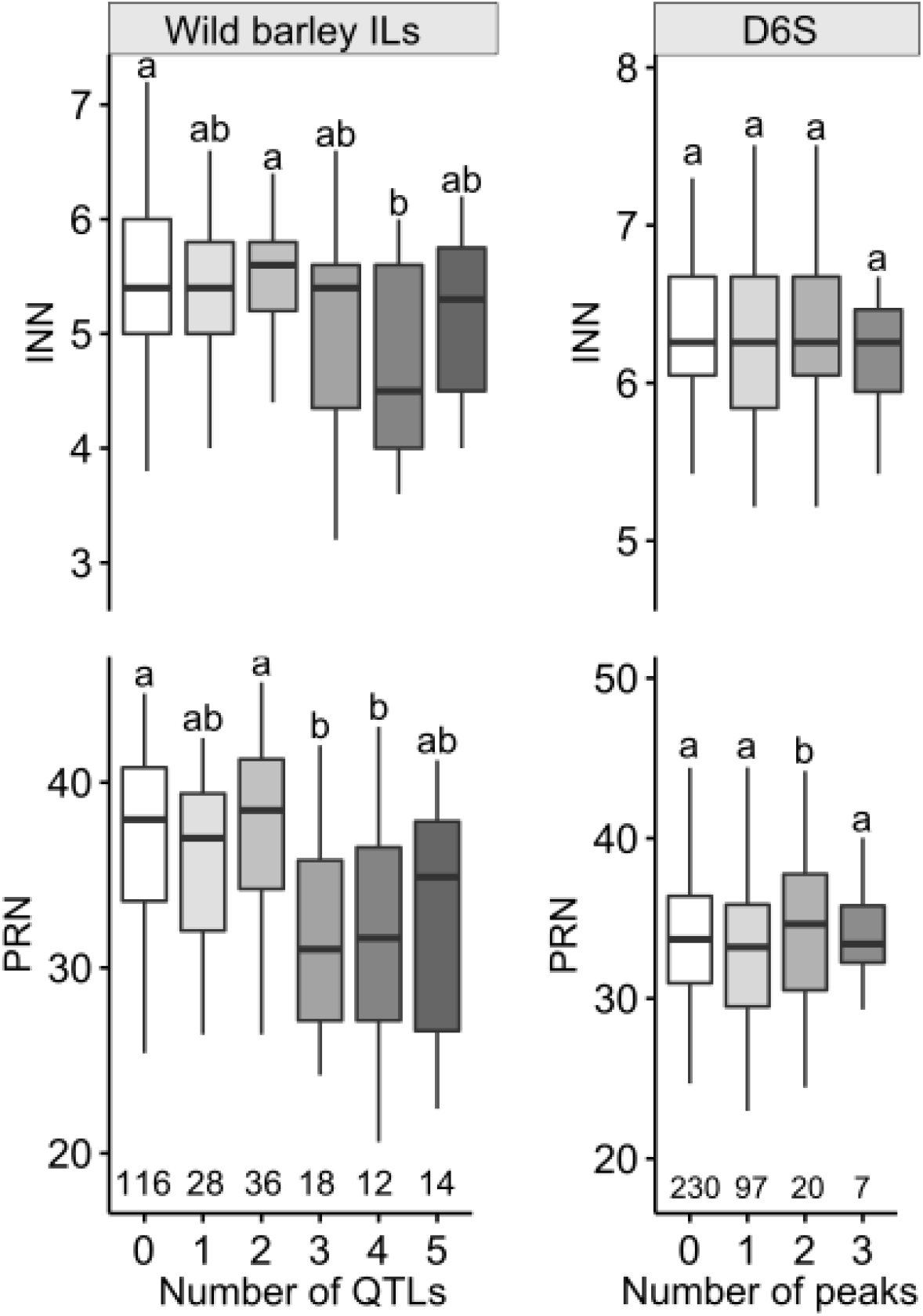
Effect of the super-locus on phytomer initiation traits. Note that similar additive effect for internode elongation was not observed for node initiation (PRN and INN). Letters above boxplot represent statistical significance from one-way analysis of variance (ANOVA) followed by Tukey–Kramer honestly significant difference (HSD) tests.

**Fig. S12.**
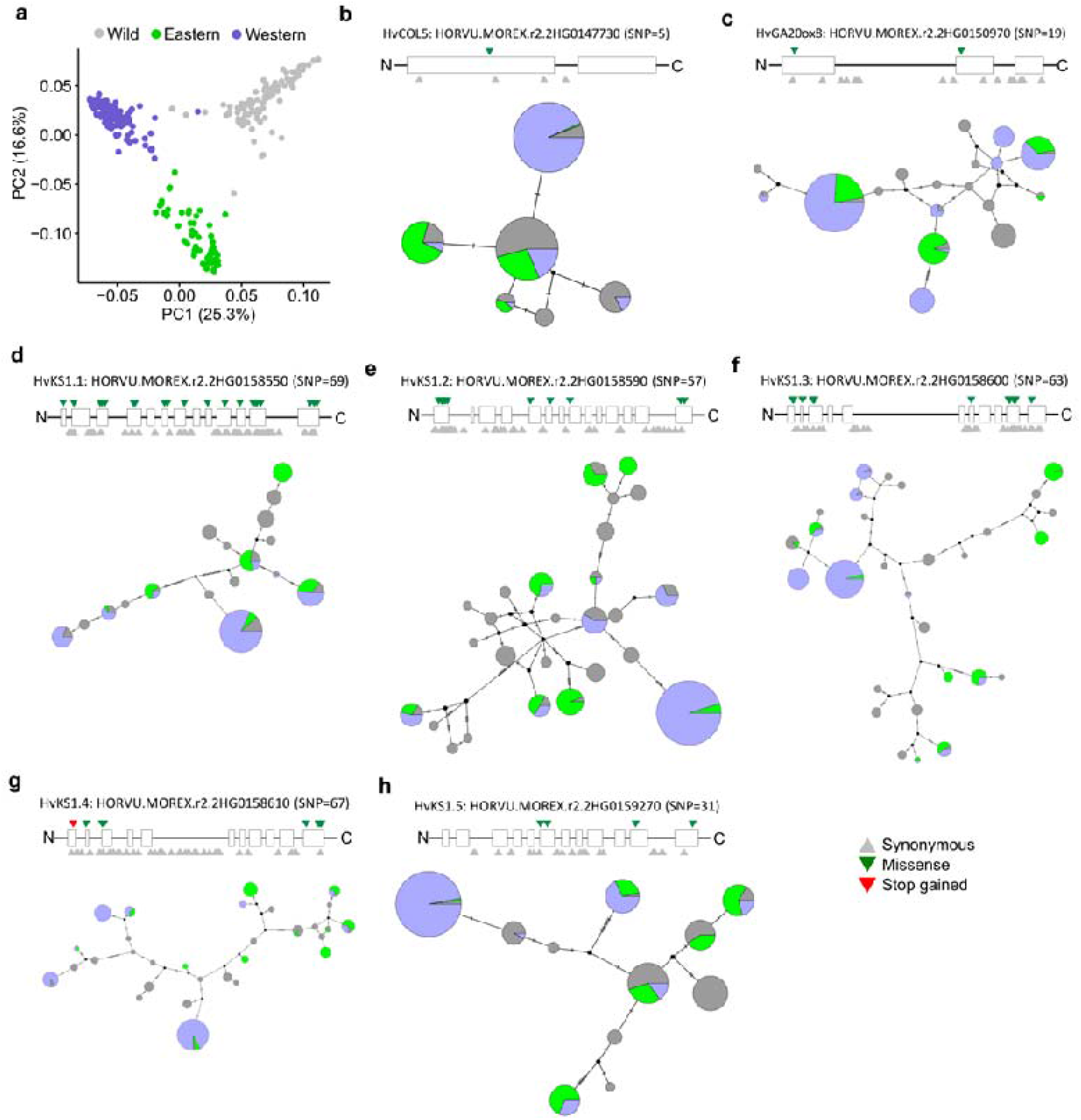
Haplotype diversity of the seven prime candidates at the superlocus. **a.** Genetic relationship of wild and world-wide domesticated barleys. A PCA based on 31,586,183 genome-wide SNPs reported previously^41^ separates domesticated barleys from the wild barleys, and further defines the domesticated barleys into two clades coincides with Eastern and Western origins. **b – h**. Haplotype diversity of the seven prime candidates. SNPs from the genomic region (from start codon until stop codon) of each gene were used to construct a median joining network. Different node colors indicate different clades defined in (**a**). Size of the nodes are proportional to the number of accessions in each node. Gene models (based on Morex v2) are shown above the networks. White boxes represent gene exons. Synonymous SNPs are indicated with grey triangles below each gene model; SNPs that induce amino acid substitutions (non-synonymous) or gain of stop codon are indicated with green or red triangles above each gene model.

**Fig. S13.**
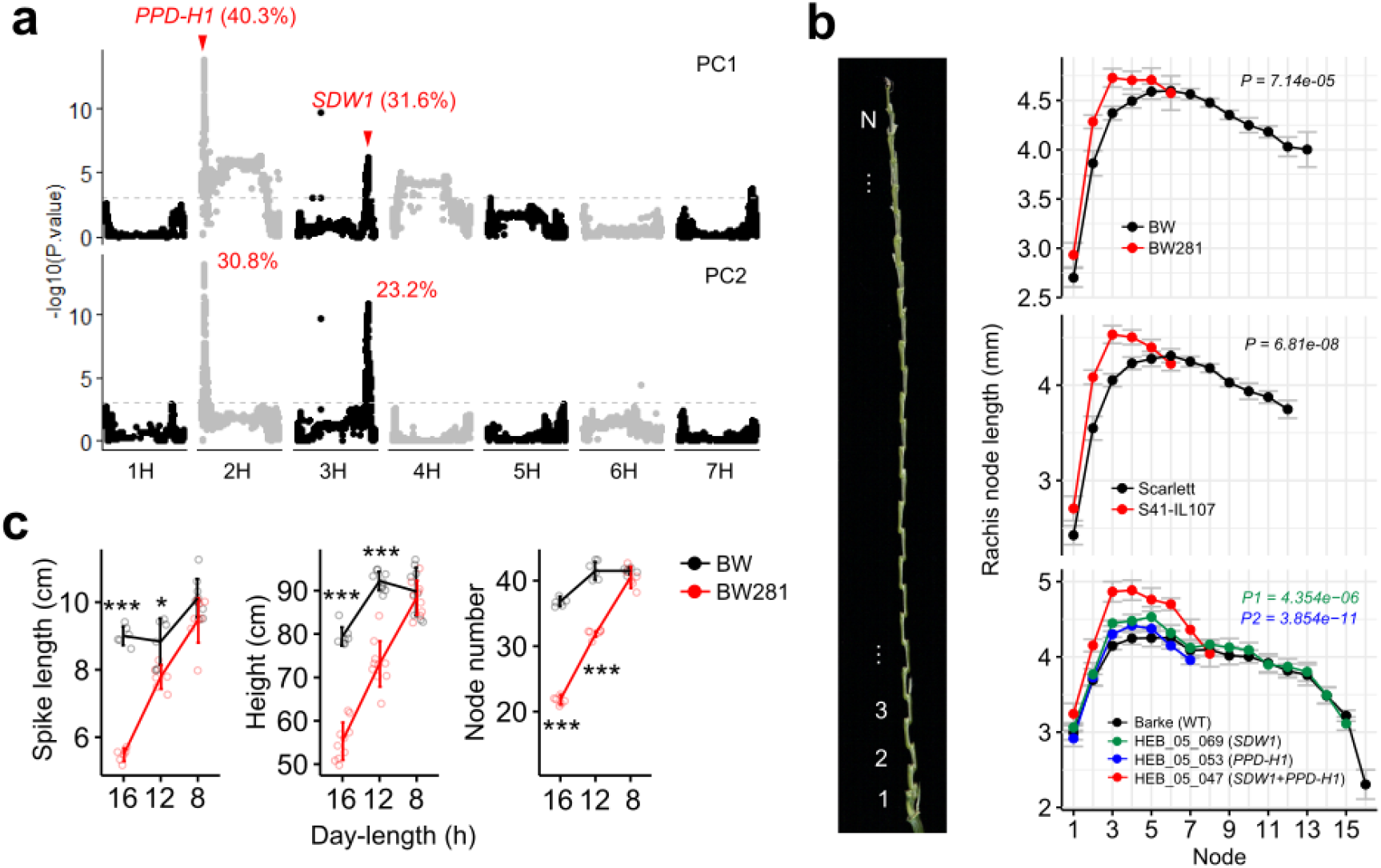
Effects of wild barley alleles at the *PPD-H1* and *SDW1* loci. **a.** Manhattan plots showing the associations of the *PPD-H1* and *SDW1* loci for the first (PC1) and second (PC2) PCA loadings based for the 14 traits. Percentage of variants explained by each loci (*R^2^*) are indicated. Grey dashed lines are genome-wide threshold at *P* = 0.001. **b.** *PPD-H1* and *SDW1* positively controls rachis internode length. Left panel illustrates the measurement of rachis internode length (one side only). Measurements were conducted on two *PPD-H1* isogenic lines and their backgrounds [BW281 vs BW (top); S42-IL107 vs Scarlett (middle)], as well as selected introgression lines from the HEB-25 with different wild barley allele introgressions (bottom). Significant values were determined by ANOVA. P1 is from the comparison between HEB-05-047 and HEB-05-069; P2 is from the comparison between HEB-05-047 and HEB-05-053. **c.** Quantitative comparison of spike length (left), plant height (middle) and potential rachis node number (right) between BW and BW281 under 16, 12 or 8 hours (h) of day-length conditions. Significant levels are determined from two-tailed Student’s *t*- test. \**P* < 0.05; \*\*\**P* < 0.001. *n* = 5 – 9 replicates.

**Fig. S14.**
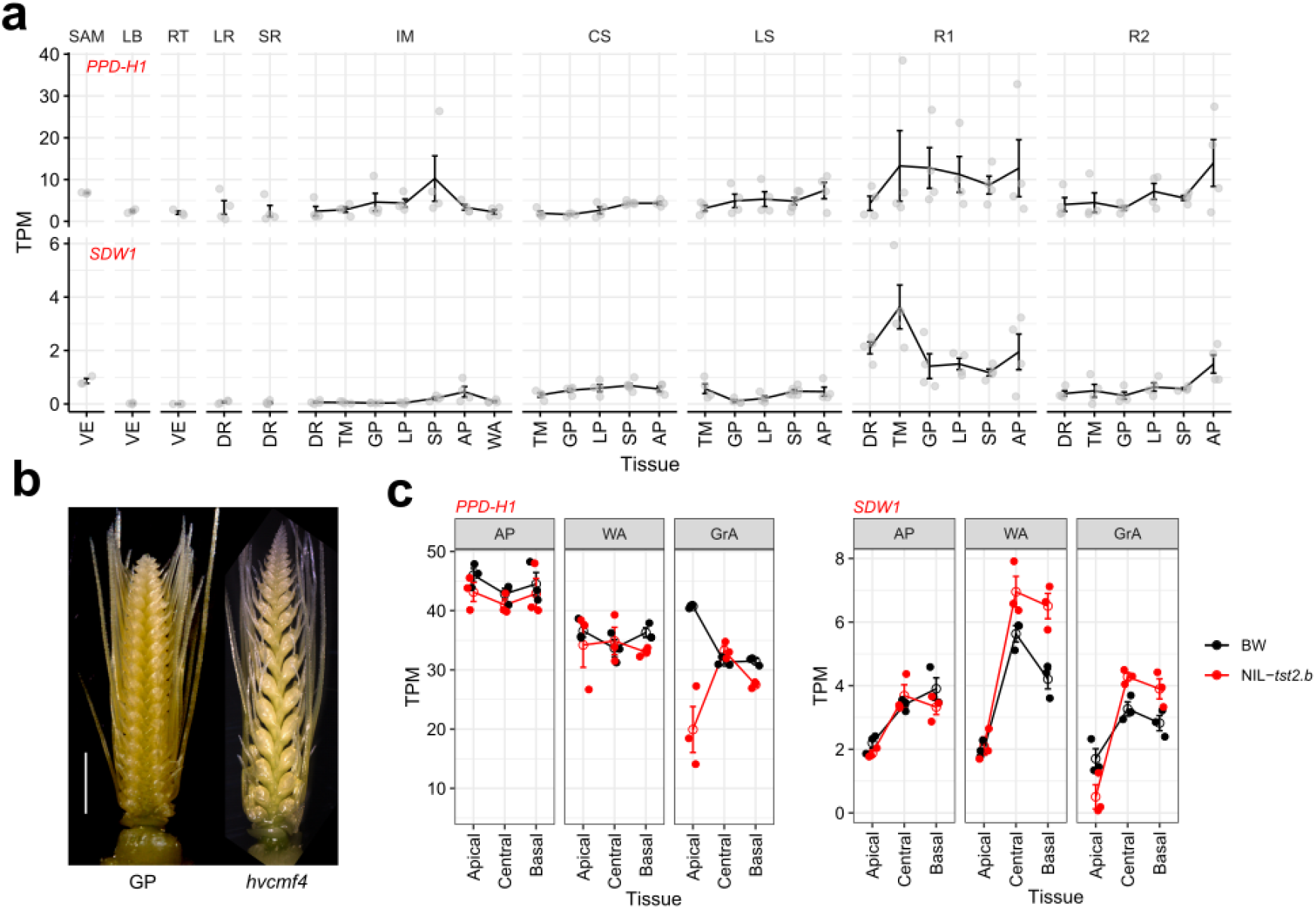
*SDW1* co-expresses with *PPD-H1* from diverse tissues. **a** – **c.** TPM values of *PPD-H1* and *SDW1* from floral meristems (SR, IM, CS, LS, R1 and R2) and non-floral tissues (SAM, LB, RT and LR) in BW (**a**), or different spike sections at three developmental stages in BW and *tst2.b* mutant (Huang *et al.*, 2023) (**b, c**). SAM, shoot apical meristem; LB, leaf blade; RT, root tips; LR and SR, leaf- and spikelet ridges; IM, inflorescence meristem; CS and LS, central and lateral spikelet; R1, rachis; R2, whole spike sections; VE, vegetative stage; DR – WA: spike developmental stages ranging from double ridge (DR), triple-mound (TM), glume primordium (GP), lemma primordium (LP), stamen primordium (SP), awn primordium (AP) and white anther (WA); GrA, green anther stage. GP, Golden Promise. Scale bar: 2mm.

**Fig. S15.**
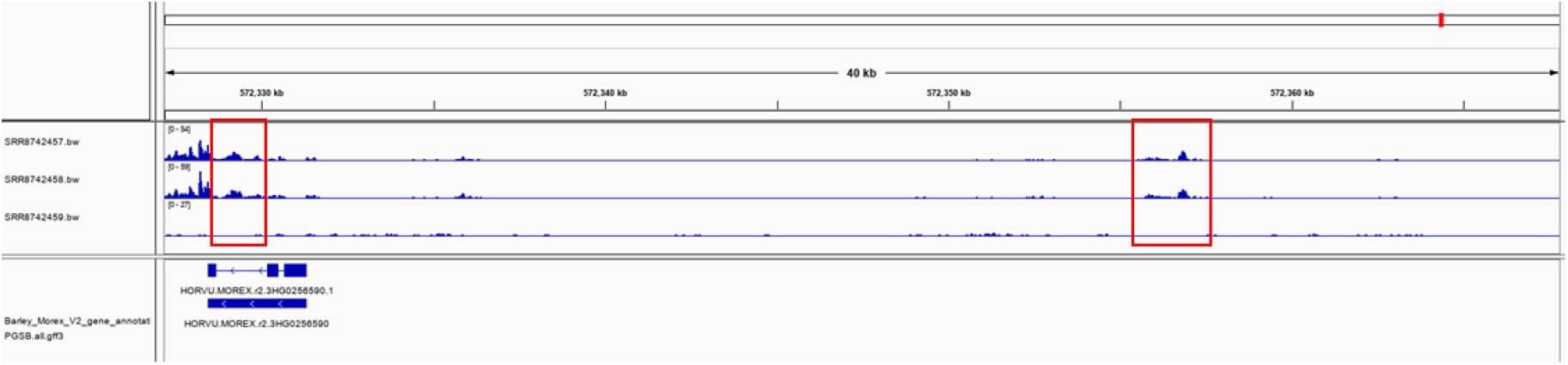
Accessible chromatin regions (ACRs) at *SDW1*. A snapshot of the Integrated Genomics Viewer browser showing the landscape of ACRs around *SDW1*. ATAC-seq data from^90^ are aligned to Morex reference v2. Identification of ACRs were done according to^90^. Track 1 and 2 are ATAC-seq reads, track 3 is a control. Red frames are two ACRs at *SDW1*, together with the 2-kb promoter regions, are used for dual-LUC assay.

**Fig. S16.**
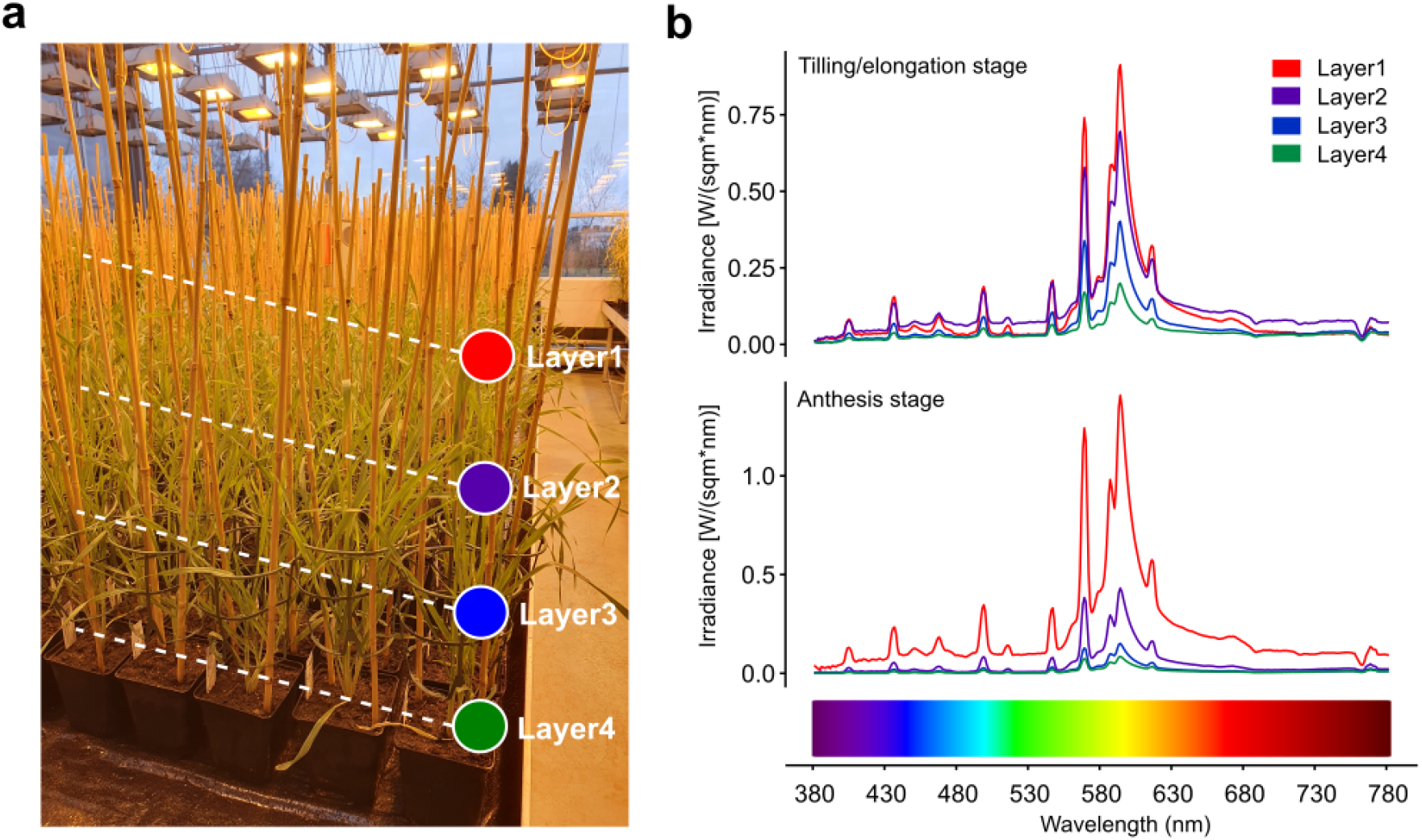
Light regimes at different barley canopy layers from the greenhouse. **a.** A representative image depicting the different canopy layers at tillering/stem elongation stage. **b.** Light regimes at different barley canopy layers during tillering/stem elongation or anthesis stages. A gradient descent of overall light intensity from Layer1 – Layer4 (distal – proximal) is observed for both stages. Note that light penetrating to the proximal end becomes severely blocked along with growth (tilling – anthesis stage).

## Other supplementary information include

Table S1. Line information and BLUE values of the phenotypic data

Table S2. Raw phenotypic data

Table S3. Summary of the QTLs detected in the wild barley population

Table S4. Summary of the QTLs detected in the D6S population

Table S5. Summary of the candidate genes identified in the D6S population

Table S6. GO enriched terms of the candidate genes identified in the D6S

Table S7. Flowering time genes orthogroups in Arabidopsis and barley

Table S8. Primers used in this study

